# CRISPR-RfxCas13d screening uncovers Bckdk as a post-translational regulator of the maternal-to-zygotic transition in teleosts

**DOI:** 10.1101/2024.05.22.595167

**Authors:** Luis Hernandez-Huertas, Ismael Moreno-Sanchez, Jesús Crespo-Cuadrado, Ana Vargas-Baco, Gabriel da Silva Pescador, José M. Santos-Pereira, Ariel A. Bazzini, Miguel A. Moreno-Mateos

## Abstract

The Maternal-to-Zygotic transition (MZT) is a reprograming process encompassing zygotic genome activation (ZGA) and the clearance of maternally-provided mRNAs. While some factors regulating MZT have been identified, there are thousands of maternal RNAs whose function has not been ascribed yet. Here, we have performed a proof-of-principle CRISPR-RfxCas13d maternal screening targeting mRNAs encoding protein kinases and phosphatases in zebrafish and identified Bckdk as a novel post-translational regulator of MZT. *Bckdk* mRNA knockdown caused epiboly defects, ZGA deregulation, H3K27ac reduction and a partial impairment of miR-430 processing. Phospho-proteomic analysis revealed that Phf10/Baf45a, a chromatin remodeling factor, is less phosphorylated upon Bckdk depletion. Further, *phf10* mRNA knockdown also altered ZGA and Phf10 constitutively phosphorylated rescued the developmental defects observed after *bckdk* mRNA depletion. Altogether, our results demonstrate the competence of CRISPR-RfxCas13d screenings to uncover new regulators of early vertebrate development and shed light on the post-translational control of MZT mediated by protein phosphorylation.

## Introduction

The first stages of metazoan development are essentially controlled by instructions deposited in the oocyte in the form of proteins and mRNA. This maternal contribution directs the first molecular events in the zygote and triggers an embryonic reprogramming where cells transition from pluripotent to differentiated cellular states^1,2^ a process named the maternal-to-zygotic transition (MZT)^2,3^. The MZT is associated with two major events. First, zygotic genome needs to be activated, and is associated with widespread chromatin remodeling within the genome^4^. Second, maternal mRNAs are selectively and actively eliminated from the embryo in a programmed manner^2,3^.

One of the most used animal models for the study of early embryonic development and MZT is zebrafish (*Danio rerio*) where the onset of zygotic genome activation (ZGA) occurs at 64-cell stage (2 hours post fertilization, hpf), leading to a major wave of transcriptional activity over the subsequent hours as the embryo progresses towards epiboly^5,6^. Some regulators of the MZT have been described and characterized in zebrafish at different levels such as transcriptional pioneer factors (i.e. Nanog, SoxB1 and Pou5f3) or chromatin remodelers controlling ZGA, and the microRNA, miR-430, and codon optimality with a major role in the clearance of the maternal RNA after ZGA^4,7–15^. In addition, translational control of maternal mRNAs and complete nuclear pore complexes maturation are other mechanisms controlling ZGA^5,12,16–18^. Despite these advances, the maternally-instructed post-translational regulatory mechanisms such as protein phosphorylation or ubiquitination modulating MZT are much less understood specially in vertebrates^2,19^ and a systematic analysis is needed.

Although maternal screenings have been performed in non-vertebrate systems^20^, maternal gene functions have remained evasive in vertebrate models largely due to the lack of suitable approaches to systematically perturb oocyte-provided RNAs that can drive early development and MZT. This is particularly important in teleosts and other aquatic vertebrate models, as RNAi technology is not effective^21–23^ and the use of morpholinos to knockdown mRNAs can trigger toxicity, off-targeting, and undesired effects such as the activation of innate immunity and cellular stress responses^24–26^. To circumvent this, we recently optimized CRISPR-RfxCas13d to induce the specific, efficient, and cost-effective degradation of RNAs in different animal embryos such as zebrafish, medaka, killifish and mouse, recapitulating well-known maternal and/or zygotic embryonic phenotypes^27,28^. This opened an unprecedented opportunity to systematically study the maternal RNA contribution that can control MZT in an early vertebrate embryo model.

Here, as a proof-of-principle, we applied our optimized CRISPR-RfxCas13d system^27,28^ to deplete 49 maternal mRNAs encoding proteins with a potential role in the regulation of protein phosphorylation during MZT. We identified seven candidates whose knockdown (KD) triggers epiboly defects, a phenotype consistent with an alteration in MZT and ZGA ^5,9,10,27,29^. By analyzing the transcriptomic profiling at the onset of the ZGA major wave, we demonstrated that the specific mRNA depletion of two kinases, Bckdk and Mknk2a, triggered a down-regulation of the pure zygotic genes (PZG); i.e. those with absent or low maternal contribution^29^. Particularly, the KD of *bckdk* mRNA, coding a kinase with a mitochondrial and cytosolic location^30–34^, led to the most severe developmental phenotype and transcriptional perturbation. Using phospho-proteomics and metabolic assays we showed that the Bckdk non-mitochondrial role was responsible for the MZT regulation that we further investigated. By performing RNA-seq later during early development, as well as SLAM-seq and ATAC-seq, we showed that ZGA and MZT were globally altered. Notably, while chromatin accessibility slightly changed upon *bckdk* mRNA KD, Histone H3K27 acetylation (H3K27ac), an epigenetic mark crucial to trigger ZGA ^5,8,35^, was significantly reduced. We also observed an impairment of the maternal RNA decay mediated by miR-430 whose biogenesis was affected. Through a phospho-proteomic analysis, we demonstrated that *bckdk* mRNA depletion reduced the phosphorylation of different proteins during MZT and we focused on Phf10/Baf45a, a protein related to chromatin remodeling that is part of the Polybromo-associated BAF (pBAF) complex belonging to the SWI/SNF family^36–38^.CRISPR–RfxCas13d knockdown of *phf10* maternal mRNA also triggered a ZGA deficiency together with epiboly defects. Importantly, Phf10 constitutively phosphorylated rescued the developmental defects observed in the absence of maternal *bckdk* mRNA. Finally, *bckdk* mRNA depletion also induced an early development perturbation and downregulation of PZG in medaka (*Oryzias latipes*), indicating a conservation of Bckdk role in MZT among teleosts. Altogether, our results i) demonstrate the competence of CRISPR-RfxCas13d as an RNA KD tool to perform maternal screenings in vertebrates, ii) uncover Bckdk as a novel post-translational modulator of MZT through the regulation of both ZGA and maternal RNA clearance and iii) demonstrate that the early developmental alterations seen after *bckdk* mRNA KD is, at least in part, due to changes in the phosphorylation state and activity of Phf10.

## Results

### A CRISPR-RfxCas13d screening for maternally-provided mRNA encoding protein phosphorylation regulators

To systematically determine whether the phosphorylation state of proteins plays a role during MZT, we selected 49 genes annotated as kinases, phosphatases and factors directly related to their activity that had not been previously characterized in zebrafish. These candidates showed a specific transcriptomic and translational pattern described for known regulatory factors of MZT such as Nanog or Pou5f3 (Oct4): highly abundant mRNA in the oocyte as well as highly translated at the onset of ZGA with subsequent decreases in translation afterwards^5,29,39,40^. Next, we designed three chemically synthesized and optimized guide RNAs (gRNAs) per mRNA to maximize targeting efficacy^28,41^ (see Methods). These gRNAs were co-injected together with a purified RfxCas13d protein with a cytosolic location^27^ into one stage zebrafish embryos (Fig. 1B, Supp. Table 1). Then, the development of these embryos was monitored for 24 hours as well as control embryos (uninjected or injected with RfxCas13d only). While most KD candidates did not affect early development compared to the control embryos, 7 KD induced epiboly defects in, at least, one third of the injected embryos, a phenotype potentially compatible with an alteration in MZT and ZGA^5,8,27^ (Fig. 1C).These 7 candidates included i) proteins related to calcium signaling that have been associated with the modulation of kinase activity, such as the Calmodulin paralogs Calm1a and Calm2a^42^ and Cab39l^43^, ii) MAPK interacting serine/threonine kinases, such as Mknk1 and Mknk2a^44,45^, and iii) a regulatory subunit part of the Serine/threonine phospho-protein phosphatase 4 complex, Ppp4r2a^46^. Interestingly, the most severe delay was observed upon the KD of the branched-chain ketoacid dehydrogenase kinase (*bckdk*) mRNA. Bckdk is a well-conserved kinase (89% and 93% amino acid similarity between human and zebrafish and between mouse and zebrafish, respectively) located in the mitochondria with a known role in controlling branched-chain essential amino acid (valine, leucine, and isoleucine, BCAA) catabolism^30,47^. In addition, Bckdk has been recently detected in the cytosol where, for example, it is able to i) upregulate the MEK-ERK signaling pathway promoting cell proliferation, invasion, and metastasis and/or ii) stimulate lipogenesis by phosphorylating ATP-citrate lyase^31–33^.

**Figure 1.**
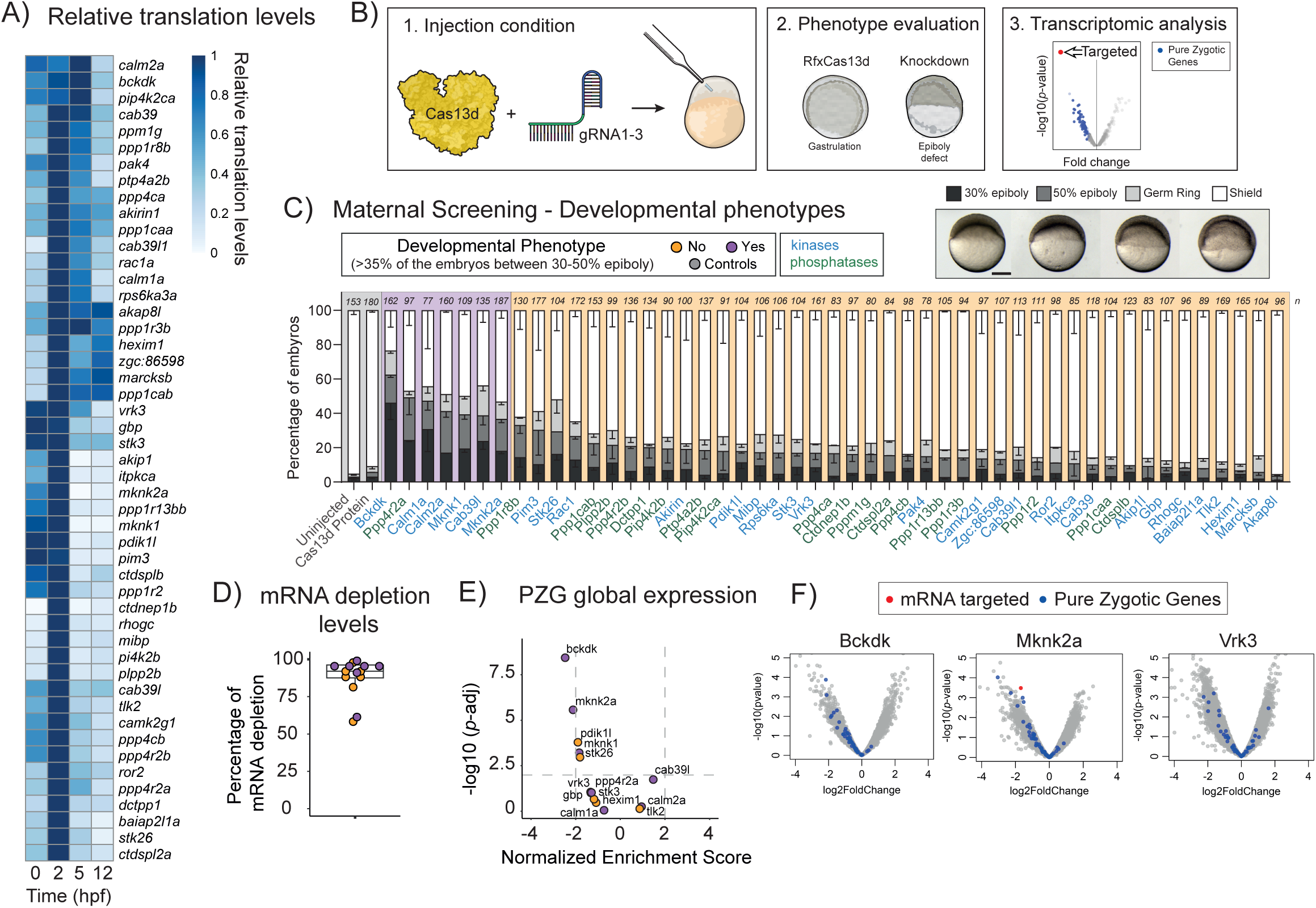
Maternal screening using CRISPR-RfxCas13d system reveals Bckdk as a potential regulator of ZGA in zebrafish. **A)** Heatmap plot of relative translation data during early developmental stages (0, 2, 5 and 12 hpf) of the kinases selected for the maternal screening. Data from Bazzini et al 2014^39^ and Chan *et al.,* 2019^5^. **B)** Schematic illustration of the experimental setup used for the maternal screening. 1. RfxCas13d protein (3ng per embryo) was mixed with a pool of 3 gRNAs (1000pg per embryo) and injected into one-cell-stage zebrafish embryos. Phenotypic evaluation at 6 hpf. Embryos injected only with RfxCas13d protein were at a normal gastrulation stage. Knockdown conditions were evaluated and considered positive when an epiboly defect was observed. 3. Transcriptomic analysis was performed at 4 hpf to evaluate mRNA depletion and the global analysis of Pure Zygotic Genes (PZG). Knockdown conditions with an epiboly defect and global downregulation of PZG were considered ZGA candidates. **C)** Stacked barplots showing the percentage of phenotypes observed at 6 hpf under maternal screening conditions. Cas13d Protein indicates control embryos injected only with RfxCas13d (3 ng/embryo). Positive candidates (purple background) were considered when more than 35% of the embryos showed a developmental delay (30-50% epiboly at 6 hpf where control embryos were at shield stage). Negative candidates and controls are indicated with orange and grey background, respectively. mRNAs encoding for kinases are labelled in blue and mRNAs encoding for phosphatases are labelled in green. The results are shown as the averages ± standard error of the mean of each developmental stage from at least two independent experiments. Number of embryos evaluated (n) is shown for each condition. Representative pictures of different zebrafish epiboly-stages from *bckdk* mRNA knockdown are shown in the upper panel. 30% epiboly, 50% epiboly, germ ring, and shield stages correspond to 4.6, 5.3, 5.7, and 6 hpf in uninjected embryos growing in standard conditions, respectively (scale bar, 0.05 mm). **D)** Boxplot with individual values showing the percentage of mRNA depletion using the RfxCas13d approach for 7 positive candidates (purple dots) and 7 negative candidates (orange dots) from maternal screening measured by Bulk-RNA-Seq at 4 hpf. The mean of mRNA depletion is represented together with the first and third quartile. Vertical lines indicate the variability outside the upper and lower quartiles. **E)** Scatter plot of Normalized Enrichment Score and adjusted *p*-value associated with Gene Set Enrichment analysis of Pure Zygotic Genes (PZG) defined by Lee *et al.,* 2013^29^ from positive and negative candidates (see methods for details). Bulk-RNA-Seq data from panel D. Vertical and horizontal dashed lines indicate -2 and 2 Normalized Enrichment Score and adjusted *p*-value = 0.01, respectively. **F)** Scatter plots representing the fold change in mRNA levels and *p-*value from two biological RNA-seq replicates (n = 20 embryos/biological replicate) at 4 hpf in zebrafish embryos injected with Cas13d protein and 3 gRNAs targeting *bckdk* (left), *mknk2a* (middle) or *vrk3* (right) mRNA. Pure Zygotic Genes mRNAs (PZG) defined by Lee *et al.,* 2013^29^ are indicated in blue. The mRNA targeted in each condition is highlighted in red.

At 24 hpf, the 7 KDs that showed an early developmental phenotype exhibited a lower viability correlating with the percentage of embryos with epiboly defects (Supp. Fig. 1A,) while the rest of the KDs did not reveal any remarkable alteration (Supp. Fig. 1B) except for Mibp, a ribosylnicotinamide kinase orthologous to human NMRK2^48^. *Mibp* mRNA KD embryos showed a defect in brain development causing microcephaly in 60% of embryos (Supp. Fig. 1C) consistent with its neural expression in zebrafish^49^ However, since we were interested in uncovering novel regulators of MZT, we focused on the 7 candidates whose KD caused an early developmental defect during the first hours after fertilization.

Epiboly failure during early zebrafish development does not always correlate with an alteration in ZGA and MZT^50^. To test whether these phenotypes were associated with a perturbation of ZGA, we performed a transcriptomic analysis of the KD of 14 candidates, including the 7 that showed an epiboly alteration and 7 with a normal development as well as embryos injected with RfxCas13d only as a control (Fig. 1B). First, RNA-seq data showed high target efficiency for the 14 targets from 58% to 99% mRNA reduction with a median of 92% (Fig. 1D). Second, to address whether ZGA was affected in the KD embryos, we compared the mRNA level of genes previously defined as pure zygotic genes^29^ (PZG), in the KD embryos to the embryos injected with Cas13d alone (Fig. 1B). Interestingly, *bckdk* and *mknk2a* KD negatively affected the expression of the PZG suggesting a role in the ZGA while other KDs such as *vrk3* did not (Fig. 1E, F and Supp. Fig. 2). Notably, other KD candidates that showed an epiboly defect did not affect ZGA but could affect other molecular pathways (Fig. 1C and 1E, Supp. Fig. 2). Indeed, analyzing downregulated genes from the 7 candidate KDs revealed that proteins related to calcium signaling and kinase activity shared the highest number of depleted genes among all possible comparisons, suggesting a potential common role for these proteins (Supp. Fig. 1F).

It has been recently shown that RNA-targeting mediated by CRISPR-RfxCas13d may trigger collateral activity in mammalian cells^51–53^. This is an uncontrolled and gRNA-independent RNA-targeting after a previous and specific gRNA-dependent activity of RfxCas13d^51–53^. One of the hallmarks of the collateral activity triggered by RfxCas13d is ribosomal fragmentation and a decrease of the RNA integrity number^51,52^. Based on these molecular markers, we did not observe evidence of collateral effects mediated by CRISPR-RfxCas13d in zebrafish embryos when targeting these 14 endogenous mRNAs (Supp. Fig. 3A-B) as we previously showed for other transcripts^27^. Altogether, these results demonstrate that i) CRISPR-RfxCas13d is a powerful KD approach to screen for dozens of maternal genes affecting early vertebrate development and ii) specifically, the KDs of two maternal mRNA encoded kinases (*bckdk* and *mknk2a)* exhibit epiboly defects together with a significant ZGA perturbation.

### *bckdk* mRNA depletion is specific, does not affect its mitochondrial activity and has a conserved developmental effect in medaka

Considering that *bckdk* mRNA KD resulted in the most drastic depletion of PZG expression and severe phenotype from the maternal screening, we focused on this kinase to further understand its role in regulating MZT. First, to validate the *bckdk* mRNA KD phenotype and rule out off-target effects, four different gRNAs were individually injected with Cas13d protein. These four gRNAs efficiently decrease *bckdk* mRNA levels (Supp. Table 1, Supp. Fig. 4A) and recapitulated the early developmental phenotype previously observed (Supp. Fig. 4B). Then, to perform a phenotype rescue experiment we used a gRNA targeting the 3’ UTR of *bckdk* mRNA which recapitulated the 6 hpf epiboly defects (Supp. Table 1, Supp. Fig. 4C and E). Despite the intrinsic mosaic nature caused by the embryo microinjection, the developmental phenotype was rescued by the coinjection of a mRNA encoding for Bckdk tagged with a HA containing a different, and gRNA resistant, 3’ UTR (Fig. 2A-B). As expected, the phenotype was not rescued by an ectopic *gfp* mRNA (Fig. 2A-B). In addition, the overexpression of *gfp* or *bckdk* mRNAs did not cause any developmental delay in WT embryos and the cognate *bckdk* mRNA was detected at mRNA and protein levels (Supp. Fig. 4D-F). Furthermore, GFP fluorescence was similar in *bckdk* mRNA KD compared to the control, reinforcing the absence of collateral activity evidence in our CRISPR-RfxCas13d targeting conditions (Supp. Fig. 4G). These results consolidate and validate the specific targeting induced by CRISPR-RfxCas13d and the role of Bckdk in regulating early development in zebrafish.

**Figure 2.**
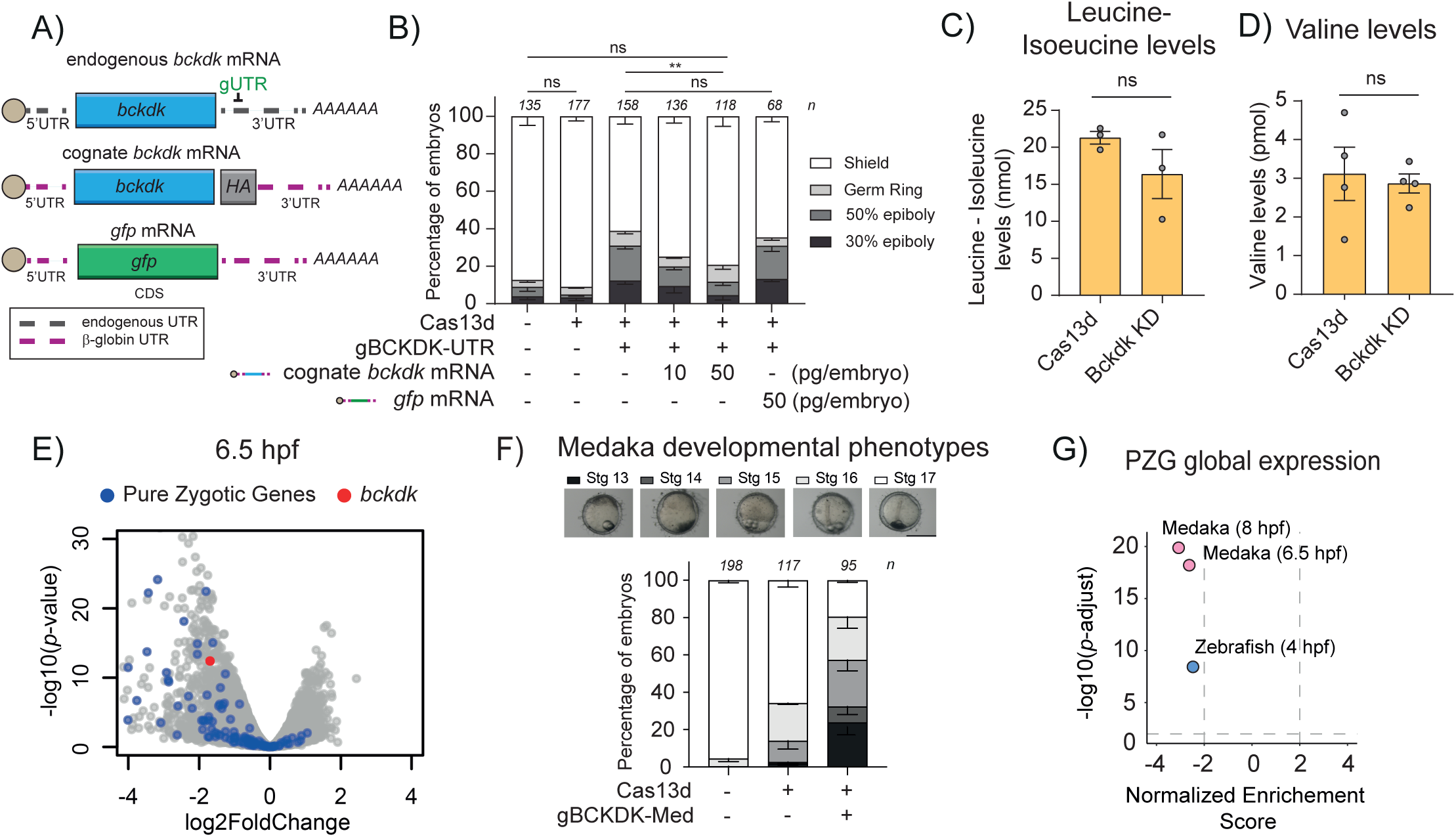
Bckdk mRNA knockdown is specific, does not affect mitochondrial function and triggers a ZGA alteration in medaka. **A)** Schematics of endogenous *bckdk*, cognate *bckdk* with β-Globin 5’ and 3’ UTR from *Xenopus laevis* and *gfp* mRNAs co-injected at 10 or 50 pg/embryo with RfxCas13d protein and one gRNA targeting the 3’UTR of the endogenous *bckdk* mRNA (gUTR). Endogenous UTR is indicated as black and dashed line and β-globin UTR is indicated as pink and dashed line. **B)** Stacked barplots showing the percentage of observed phenotypes in zebrafish embryos injected with RfxCas13d protein (3 ng/embryo) together with one gRNA targeting the 3’UTR of the endogenous *bckdk* mRNA (gBCKDK-UTR) (1000 pg/embryo) and co-injected with an exogenous *bckdk* mRNA (10-50 pg/embryo) or with a *gfp* mRNA (50 pg/embryo). The results are shown as the averages ± standard error of the mean of each developmental stage from at least two independent experiments. Number of embryos evaluated (n) is shown for each condition. The phenotype selection criteria were the same as in Fig. 1C. (ns = non-significant, \*\**p*<0.01, χ2-test). Leucine/isoleucine **(C)** and valine **(D)** normalized levels in zebrafish embryos injected with RfxCas13d alone (Cas13d) or together with a mix of 2 gRNAs targeting *bckdk* mRNA (Bckdk KD). The results are shown as the averages ± standard error of the mean of at least 3 biological replicates (n = 50 embryos/biological replicate) (ns = non-significant, Mann–Whitney U-test) **E)** Scatter plot representing the fold change in mRNA level and the associated *p-*value from three biological RNA-seq replicates (n= 10 embryos/biological replicate) at Stage 11 (Late Blastula Stage; 6.5 hpf) in medaka embryos in the conditions described in panel G. *bckdk* mRNA is represented in red. Pure Zygotic Genes mRNAs from medaka embryos determined from Li *et al.,* 2020^57^ data are depicted in blue. **F)** Stacked barplots showing percentage of observed phenotypes in medaka embryos injected with RfxCas13d protein (6 ng/embryo) alone or together with one gRNA (gBCKDK-Med) targeting *bckdk* mRNA (2000 pg/embryo). The results are shown as the averages ± standard error of the mean of each developmental stage from three independent experiments. Number of embryos evaluated (n) is shown for each condition. Representative pictures of medaka epiboly-stages from *bckdk* mRNA knockdown are shown in the top panel. Epiboly-stages earlier than Stg 16 at 25 hpf indicate epiboly defects. Stages 13, 14, 15, 16 and 17 correspond to 13, 15, 17.5, 21 and 25 hpf in uninjected embryos growing in standard conditions, respectively (scale bar, 0.1 mm). **G)** Scatter plot of Normalized Enrichment Score and adjusted *p*-value associated with Gene Set Enrichment Analysis of Pure Zygotic Genes (PZG) defined by Lee *et al.,* 2013^29^ or by Li *et al.*, 2020^57^ upon the depletion of *bckdk* mRNA in zebrafish (blue dot, 4 hpf) or medaka embryos (pink dots, 6.5 or 8 hpf), respectively (see methods for details).

Second, to determine the onset of the observed developmental defects, cell proliferation (mitotic cells) and total number of cells were quantified in embryos depleted on Bckdk before the beginning of the epiboly at 2 and 4 hpf (Supp. Fig. 4 H-J). No significant differences were observed between embryos with KD of *bckdk* mRNA and the control condition, suggesting that Bdkck depletion did not affect the proliferation rate nor cell division and demonstrating that the observed developmental phenotype takes place between 4 and 6 hpf (Supp. Fig. 4H-J).

Third, the loss-of-function of Bckdk in mammalian cells causes a loss of phosphorylation and hyperactivation of the branched-chain ketoacid dehydrogenase Bckdh in the mitochondria^30^ resulting in a decreased BCAA availability^54–56^. To elucidate whether the KD of *bckdk* mRNA affects its mitochondrial activity during early development, we measured BCAA levels. We observed that the amount of isoleucine-leucine and valine did not significantly change in comparison to embryos injected with Cas13d alone (Fig. 2C and D). Together, these data suggest that the *bckdk* mRNA KD only affects its non-mitochondrial function ultimately triggering a ZGA alteration and epiboly defects.

Fourth, *bkcdk* mRNA is also maternally provided in medaka^57^, another teleost model used to study MZT and that evolutionary diverged from zebrafish 115-200 millions of years ago^58^. To test whether the role of Bckdk is conserved among teleosts, the maternal *bckdk* mRNA in medaka was depleted using CRISPR-RfxCas13d (Supp. Table 1). As observed in zebrafish, CRISPR-RfxCas13d induced an efficient KD of *bckdk* mRNA (3.2-fold change p < 3.3e-13) at 6.5 hpf in medaka (Fig. 2E). Additionally, the downregulation of *bckdk* mRNA caused i) a delay during early development (Fig. 2G), ii) a decrease in viability later during the embryogenesis (Supp. Fig. 4K) and iii), based on RNA-seq, triggered a global downregulation of PZG observed at different time points along the ZGA in medaka^57^ (Fig. 2E and G, and Supp. Fig 4L). Therefore, the role of Bckdk during embryogenesis is conserved between zebrafish and medaka.

### Bckdk modulates the transcription of zygotic and maternal-zygotic expressed genes

Upon determining that *bckdk* mRNA KD impacts the PZG expression, we explored whether it also altered the transcription of the maternal-and-zygotic expressed genes which represent more than half of the transcriptome^6,59,60^. We employed SLAM-seq, a technique that differentiates maternal and zygotic mRNA through nascent mRNA labeling with 4-thiouridine (S4U) (Fig. 3A). To accurately assess these effects, we first optimized SLAM-seq alongside CRISPR-RfxCas13d use in zebrafish. Thus, we titrated different amounts of S4U that in combination with RfxCas13 protein did not affect early development. Previously, it has been showed that up to 75 pmol of S4U per embryo resulted in high nascent label signal without developmental alteration^6,59^. Although injection of *cas13d* mRNA with 75 pmol of S4U did not affect early embryogenesis, 75 pmol or 50 pmol of S4U injected in combination with purified RfxCas13d protein caused epiboly defects (Supp. Fig. 5A). For that reason, we injected 25 pmol of S4U per embryo to address whether, beyond PZG, Bckdk depletion affected the transcription of the maternal-and-zygotic genes (Fig. 3A and Supp. Fig. 5B). As expected and similar to the previous RNA-seq experiment (Fig. 1E-F and Supp. Fig. 2, *bckdk* panel), *bckdk* mRNA and PZG were downregulated in the KD embryos compared to the control embryos (Supp. Fig. 5B and C). Beyond previously described PZG^29^, we observed a downregulation of an additional set of PZG (Fig. 3B and C, Supp. Fig. 5D) recently defined through SLAM-Seq approaches^6^. Interestingly, besides the PZG, 1734 genes with maternal mRNA contribution, displayed a reduction in labeled transcript levels in the KD compared to the control, indicating that Bckdk also affects the transcription of maternal-zygotic genes. In total, 1884 genes were less transcribed upon *bckdk* mRNA KD (2-fold change) denoting a global ZGA downregulation (Fig. 3B). Nevertheless, there were 2907 genes which zygotic expression (label reads) was not affected in the *bckdk* mRNA depletion, suggesting that their transcription was independent of Bckdk (Fig. 3B), and 1491 mRNAs were upregulated (Fig. 3B). The vast majority of these genes (96.4%) were normally transcribed between 4 and 7 hpf in WT embryos^6^ (Fig. Supp Fig. 5E). Notably, 36.4% of the genes up-regulated in *bckdk* mRNA KD at 4 hpf did not show any transcriptional signal, based on SLAM-seq analysis, in WT embryos at that time point^6^ suggesting that Bckdk depletion could trigger an acceleration of genome activation for a subset of genes (Supp. Fig. 5F). These data suggest that *bckdk* mRNA KD affects the transcription dynamics of genes specifically expressed during ZGA and not genes activated later in zebrafish development. Remarkably, down-regulated genes showed enriched biological processes related to transcriptional regulation and RNA processing meanwhile up-regulated genes revealed processes associated with translation and protein biogenesis (Supp. Fig. 5G). In conclusion, our results show that SLAM-seq in combination with CRISPR-Cas13d KD can be used to dissect the impact of a maternal factor in transcription during the MZT, and, specifically, that *bckdk* mRNA depletion deregulates the transcription dynamics of a more than half (54%) of the genes expressed during ZGA in zebrafish.

**Figure 3.**
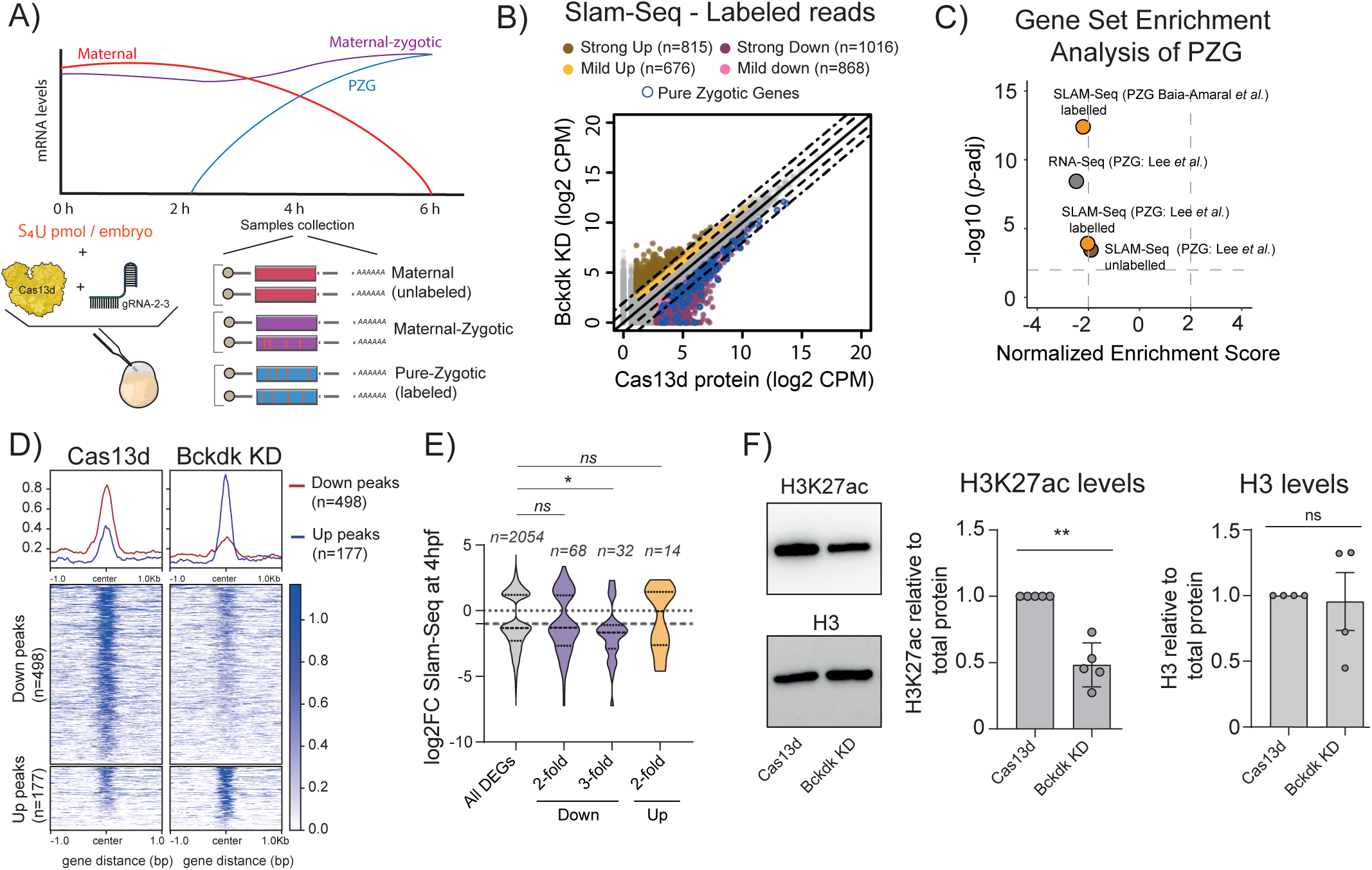
The knockdown of *bckdk* mRNA alters gene expression, slightly affecting chromatin accessibility, and significantly impacting H3K27ac levels. **A)** Schematic representation of SLAM-Seq workflow. Zebrafish embryos were injected with 4-thiouridine (S_4_U, 25pmol/embryo) in one-cell-stage together with RfxCas13d protein alone or with a mix of 2 gRNAs targeting *bckdk* mRNA. Total RNA was extracted at 4 hpf from 20 embryos. Total RNA was chemically modified before library preparation. After library preparation T>C ratio conversion was used to differentiate between unlabelled (pre-existing RNA; Maternal) and labelled (newly synthesized RNA; Zygotic) reads (see Methods for further details). **B)** Scatter plot showing newly transcribed RNAs (SLAM-seq data in Counts per million reads; CPM) of zebrafish embryos at 4 hpf from 2 and 4 biological replicates (n= 25 embryos/biological replicate) from embryos injected with RfxCas13d protein alone (Cas13d protein) or with a mix of 2 gRNAs targeting *bckdk* mRNA (Bckdk KD), respectively. Dashed and dot-dash lines indicate a 2 and 5-fold difference between RNA levels, respectively. Up- and down-regulated genes are represented by yellow (between 2 and 5-fold change) and brown (> 5-fold change) or pink (2 and 5-fold change) and purple (> 5-fold change) dots, respectively. The number of mRNAs in each category is shown (n). **C)** Scatter plot of Normalized Enrichment Score and adjusted *p*-value associated with Gene Set Enrichment analysis of Pure Zygotic Genes (PZG) defined by Lee *et al.,* 2013^29^ (PZG: Lee et al.,) or Baia-Amaral et al., 2024^6^ (PZG: Baia-Amaral et al.,) from Bckdk KD data generated by RNA-Seq (grey dot) or by SLAM-Seq with labelled (orange dots) or unlabelled (brown dot) reads (see methods for details). **D)** Plot profile (top) or heatmaps (bottom) representing normalized ATAC intensity signal for less accessible regions (red line; Down Peaks) and more accessible regions (blue line; Up peaks) from the comparison between 2 biological replicates (n= 80 embryos/biological replicate) from zebrafish embryos injected with RfxCas13d protein (Cas13d) alone or together with a mix of 2 gRNAs targeting *bckdk* mRNA (Bckdk KD). Number of less accessible (Down peaks) or more accessible (Up peaks) regions are shown (n). **E)** Violin plots showing the distribution of Log2 fold change of RNA levels (SLAM-Seq) from all differential expression genes (DEGs) and those associated with less accessible (Down) or more accessible (Up) regions from ATAC-Seq data. Dash lines and dot lines inside the violin plots indicate the mean and quartiles, respectively. Grey dot line and dash line outside the violin plots indicate 0-fold and 1.5-fold in RNA levels, respectively. (ns=non-significant, \**p*<0.05, Mann– Whitney *U*-test). Number of differential expression genes (n) for each category are shown. **F)** Representative western blot images for H3K27ac and H3 of embryos injected with RfxCas13d protein alone (Cas13d) or together with 2 gRNAs targeting *bckdk* mRNA (Bckdk KD) (left). Barplots represent H3K27ac or H3 levels relative to total proteins as the averages ± standard error of the mean at least four biological replicates from two or three independent experiments. Zebrafish embryos were collected at 4 hpf (n= 25 embryos/biological replicate) (\**p*<0.05, one sample t-test) (Right).

### *bckdk* mRNA KD leads to a slight rewiring of chromatin accessibility but a strong reduction of H3K27ac during MZT

To address whether the transcriptional changes were associated to altered chromatin accessibility, we performed Assay for Transposase-Accessible Chromatin analyses coupled to high-throughput sequencing (ATAC-seq) in *bckdk* mRNA KD and WT (Cas13d injected embryos) at 4 hpf. Interestingly, we observed that *bckdk* mRNA depletion only induced a slight change in chromatin accessibility with hundreds of regions with differential openness (n=675; ∼3% of the total peaks), compared to embryos only injected with Cas13d (Fig. 3D and Supp. Fig. 5H). Most of these regions (n=498, 74% of differentially accessible regions) were less accessible upon *bckdk* KD and, to some degree, enriched in some particular transcription factor motifs, including Gata1/2/3, Foxo6, Six5 and CTCFL (Supp. Fig. 5I). On the other hand, more accessible regions (n=177, 26% of differentially accessible regions) showed a minor enrichment in different transcription factor motifs, such as Nf1, Tfcp2l1, Hltf, Nfkb1 and Tead4 (Supp. Fig. 5J). Overall, the motif analyses showed relative low enrichments compared with the loss-of-function of specific transcription factors^8,10,61,62^, suggesting a modest and less specific effect of *bckdk* mRNA KD on chromatin accessibility that likely affects more heterogenous transcription factor binding sites. Differentially expressed genes (DEG) according to Slam-seq labeled data (Fig. 3B) associated with regions altered in their openness did not reveal a strong correlation towards transcriptional miss-regulation in the same direction as genome accessibility changes. Nevertheless, we found that downregulated genes showed a minor but still significant difference towards less open chromatin regions (Fig. 3E less accessible regions >3-fold change, *p*-value = 0.0415, Mann–Whitney U-test).

Since this slight impact on chromatin accessibility, we hypothesized that other layer of transcriptional regulation could be affected upon Bckdk depletion. Indeed, it has been shown that chromatin accessibility is an initial step, induced by pioneer factors, to activate transcription after fertilization but is not the ultimate trigger of ZGA^8^. Instead, the inhibition of Histone acetylation caused a massive downregulation of ZGA^5^ without altering the chromatin accessibility landscape^8^. Indeed, we observed a prominent and significant decreased in H3K27ac (*p*-value = 0.0023, Welch’s t-test), an epigenetic mark essential to promote ZGA, in *bckdk* mRNA KD embryos in comparison to control embryos without changes in total H3 amount (Fig. 3F and G, Supp. Fig. 5K and L).

Altogether, these data indicate that *bckdk* mRNA KD leads to a slight alteration of chromatin accessibility and a strong reduction in H3K27ac during early development that might be due to the miss-regulation of global chromatin remodelers rather than specific transcriptional activators or pioneer factors.

### *bckdk* mRNA depletion affects miR-430 processing and activity during MZT

The maternal RNA degradation during the MZT is primarily driven by both maternal and zygotic encoded mechanisms^11^. Within the zygotic mechanisms, miR-430 stands out as a critical microRNA (miRNA) facilitating the clearance of maternal RNAs^11–13^. Since ZGA was notably affected upon *bckdk* mRNA KD, we wondered whether the maternal mRNA decay was also compromised. To address this, we performed a transcriptomic analysis at 6 hpf when the vast majority of maternal RNAs have been totally or partially eliminated^11,40^. While the maternal mRNAs degraded by the maternal program did not show a substantial difference in their stability when comparing *bckdk* mRNA KD to embryos only injected with RfxCas13d (D, maximal vertical distance between the compared distribution= 0.148, *p*-value= 6.525e-07, Kolmogorov-Smirnov test), those eliminated by the zygotic program were more stable and displayed higher mRNA levels (D=0.248*, p*-value < 2.2e-16, Kolmogorov-Smirnov test) (Fig. 4A). More strikingly, maternal RNAs whose clearance depended on miR-430^40^ were notably higher in the KD compared to the control embryos (D= 0.332 *p*-value = 3.102e-11; golden targets: D= 0.368 *p*-value = 3.855e-12, Kolmogorov-Smirnov test) (Fig. 4A). Therefore, these results mimic a partial miR-430 lack-of-function molecular phenotype^13^. To address whether the expression or processing of miR-430 was affected upon *bckdk* mRNA KD, we quantified the primary (pri-miR-430) and mature miR-430 by qRT-PCR in comparison to control embryos injected with RfxCas13d protein. We observed that the level of pri-miR-430 was not significant different in both control and Bckdk KD embryos at Dome or Shield stage (Fig. 4B, Supp. Fig 6A). Interestingly, the mature miR-430 levels were lower at dome stage in the *bckdk* mRNA KD compared to the control embryos suggesting an alteration in the miR-430 processing. However, the amount of mature miR-430 was recovered at shield stage (Fig. 4C and Supp. Fig. 6B). Notably, the injection of mature miR-430 in the *bckdk* KD embryos partially restored the levels of well-known mRNA targets of this miRNA that were less degraded upon *bckdk* mRNA KD (Fig. 4D) suggesting that the delayed processing of miR-430 was partially responsible of the reduced mRNA clearance.

**Figure 4.**
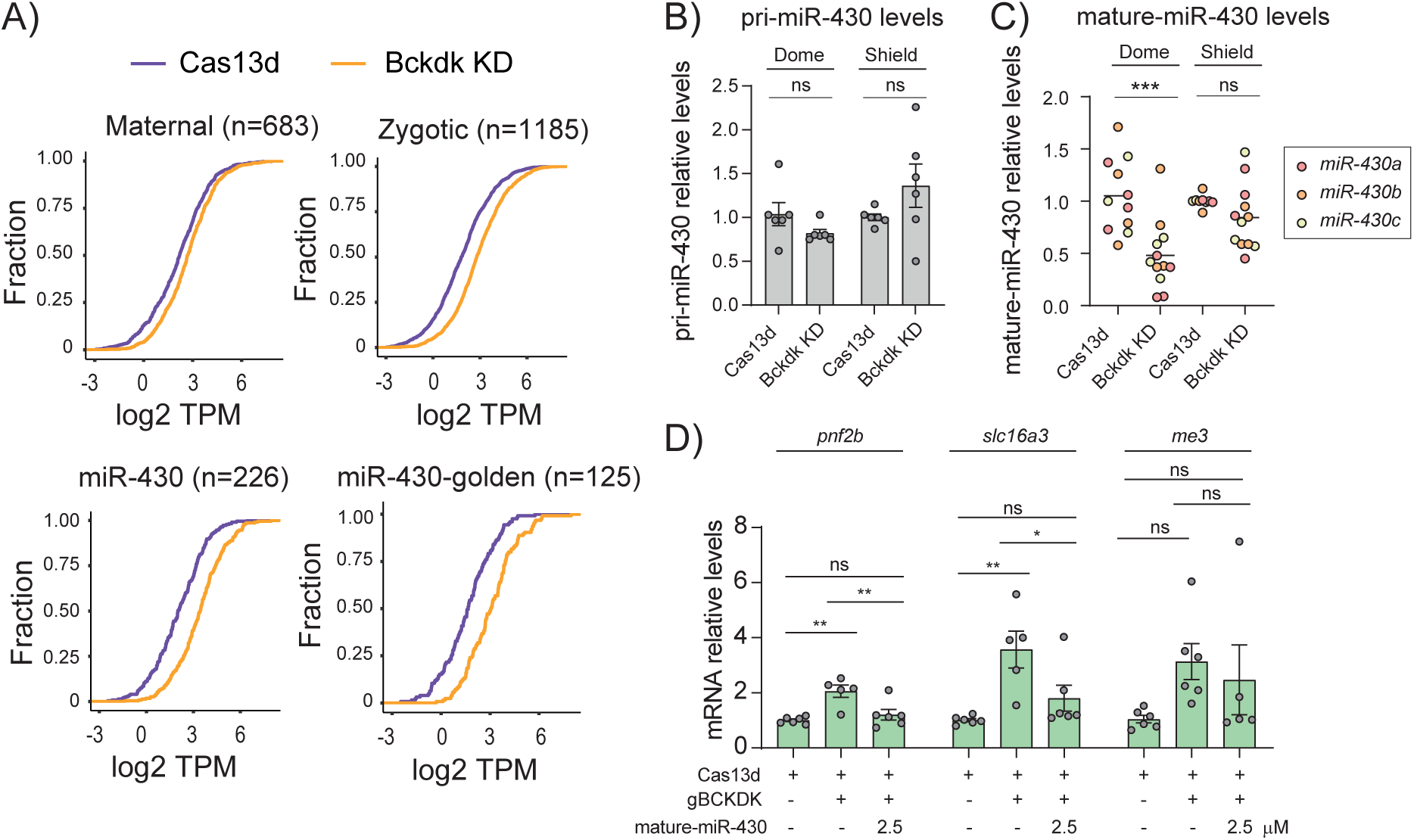
miR-430 processing and activity is compromised upon *bckdk* mRNA depletion. **A)** Cumulative distribution of mRNA levels (TPM; Transcript per million reads) in control (Cas13d; purple line) and *bckdk* mRNA knockdown embryos (Bckdk KD; orange line) that are degraded by 4 different pathways determined by Vejnar *et al.*, 2019^11^ (maternal, zygotic, miR-430) and described by Medina-Muñoz *et al.*, 2021^40^ (miR-430 golden candidates were defined as downregulated mRNAs, 4-fold, from the comparison between WT and maternal/zygotic Dicer mutant embryos at 6 hpf. Number of mRNA controlled by each pathway is shown (n). **B)** RT-qPCR analysis showing relative primary miR-430 (pri-miR-430) transcript levels at 4.3 hpf (Dome) and 6 hpf (Shield). Results are shown as the averages ± standard error of the mean from 3 experiments with 2 biological replicates per experiment (n= 10 embryos/biological replicate) for RfxCas13d protein alone (Cas13d) and RfxCas13d plus 2 gRNAs targeting *bckdk* mRNA (Bckdk KD), respectively. *taf15* mRNA was used as normalization control (ns = non-significant, unpaired t-test). **C)** RT-qPCR analysis showing levels of mature miR-430 isoforms (miR430-a, red; miR430-b, orange; and miR430c, yellow) at 4.3 hpf (Dome) and 6 hpf (Shield). Results are shown as individual values and the mean from 3 experiments with 2 biological replicates per experiment (n= 10 embryos/biological replicate) for RfxCas13d protein alone (Cas13d) and RfxCas13d plus 2 gRNAs targeting *bckdk* mRNA (Bckdk KD). ncRNA *u4atac* was used as normalization control (ns = non-significant, ***p<0.001, unpaired t-test). **D)** RT-qPCR analysis showing levels of 3 miR-430 targets that are highly stable under Bckdk KD conditions (*pnf2b*, *slc16a3* and *me3*) at 6 hpf. Results are shown as the averages ± standard error of the mean from 3 experiments with 2 biological replicates per experiment (n= 10 embryos/biological replicate). *taf15* was used as normalization control. Grubbs’s test was performed first for outlier’s identification. (ns = non-significant, *p<0.05, **p<0.01, one-way ANOVA).

Together, these results reveal that the lack of non-mitochondrial fraction of Bckdk i) in part compromises the maternal RNA decay triggered by the zygotic program and ii), more remarkably, delayed the processing of miR-430 and decreased the mRNA decay activity of this miRNA through the MZT.

### Bckdk regulates phospho-proteome during MZT and modulates Phf10 phosphorylation to control early development

Since Bckdk is a kinase, we asked whether the depletion of *bckdk* mRNA could affect the phosphorylation of proteins potentially involved in the regulation of MZT. To assess this, we performed a quantitative phospho-proteomic analysis in control embryos injected with Cas13d alone and in *bckdk* mRNA KD embryos at 4 hpf. First, 3336 phosphorylated proteins were detected in both phospho-proteomes and we observed 19 phospho-peptides from 16 proteins to be differentially phosphorylated in the *bckdk* mRNA KD compared to the control embryos (Fig. 5A; Phosphoproteins in control and KD embryos). 63% of the phosphopeptides, belonging to 9 proteins, were significantly less phosphorylated upon Bckdk KD (2-fold change p < 0.05, Fig. 5A, Supp. Fig 7A, Supp. Table 2). Notably, none of these 9 proteins are predicted to localize in the mitochondria (Supp. Fig. 7A), in contrast to a phospho-proteome upon Bckdk inhibition in rat liver where more than 55% of the identified targets localized in the mitochondria^32^. Additionally, we observed that the phosphorylation state of the Bckdha motif (tyrighhS*tsddssa) described as the target of Bckdk in the mitochondria, did not significantly changed upon *bckdk* mRNA KD (Fig. 5A, blue dots). These results strengthened the hypothesis that the early developmental phenotype and MZT perturbation in the zebrafish embryos depleted of *bckdk* mRNA was due to the non-mitochondrial activity of this kinase (Supp. Fig. 7A). Moreover, we also found 7 phospho-peptides from 7 proteins more phosphorylated (Fig. 5A, Supp. Fig. 7B, Supp. Table 2) that may be an indirect effect triggered by the absence of Bckdk. Furthermore, 11 proteins were differentially accumulated upon *bckdk* mRNA KD where 9 were down-regulated and 2 more abundant (2-fold change p < 0.05, Supp. Fig 7C, Supp. Table 2). None of the mRNAs encoding these proteins showed a significant changed by Slam-seq or RNA-seq data (Fig. 3A and Supp. Fig. 2) and 3 out the 11 proteins were also detected in the phospho-proteome without substantial differences between control and *bckdk* mRNA KD embryos. Therefore, this differential protein accumulation upon Bckdk depletion was likely due to translational or an indirect post-translational regulation.

**Figure 5.**
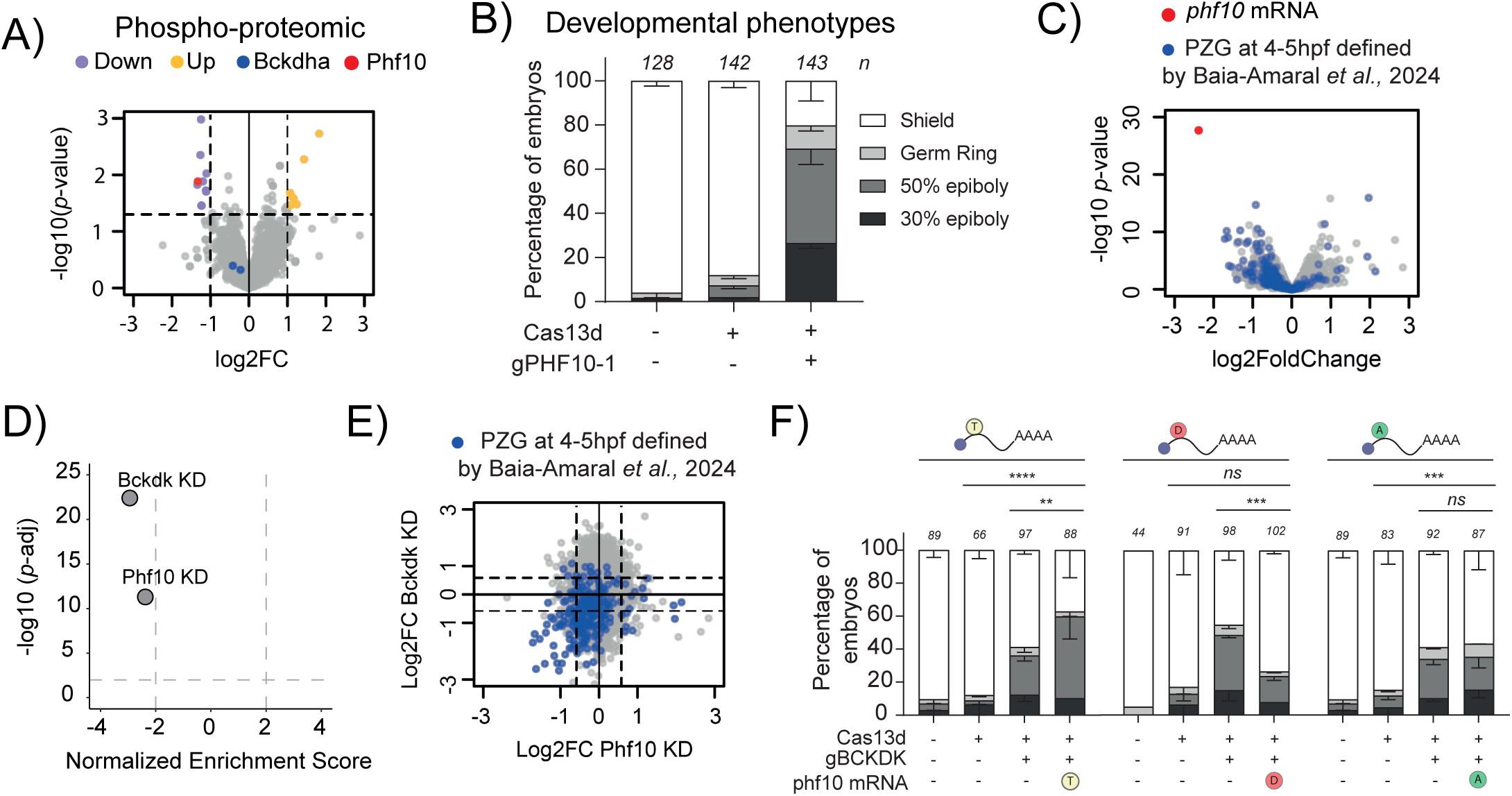
Phosphoproteomic analysis in Bckdk KD embryos uncovers Phf10 as a new regulator of ZGA. **A)** Scatter plot representing the fold change in phosphopeptides abundance levels and the associated *p*-value from 4 biological replicates (n= 100 embryos/biological replicate) at 4 hpf in zebrafish embryos injected with RfxCas13d protein alone or with a mix of 2 gRNAs targeting *bckdk* mRNA. Phosphopeptides abundance levels of Bckdha are depicted in blue. Phf10 phosphopeptide levels is represented in red. Less (Down) and more (Up) phosphorylated peptides are indicated in purple and yellow, respectively. Dashed lines indicated 2-fold in phosphorylated peptides levels and *p*-value= 0.05. **B)** Stacked barplots showing the percentage of phenotypes observed under the knockdown condition of *phf10* mRNA (injecting 3ng/embryo of RfxCas13d protein and 1000 pg/embryo of a gRNA targeting *phf10* – gPHF10-1). The results are shown as the averages ± standard error of the mean of each developmental stage from 4 independent experiments. The phenotype selection criteria were the same as described in Fig. 1C. Number of embryos evaluated (n) for each condition is shown. **C)** Scatter plot representing the fold change in mRNA level and the associated *p-*value from 3 biological replicates from RNA-Seq data (n= 20 embryos/biological replicate) at 4 hpf in zebrafish embryos injected with RfxCas13d protein alone or together with one gRNA targeting *phf10* mRNA. *phf10* mRNA is represented in red. Pure Zygotic Genes (PZG) mRNAs determined from Baia-Amaral *et al.*, 2024^6^ are shown in blue. **D)** Scatter plot of Normalized Enrichment Score and adjusted *p*-value associated with Gene Set Enrichment analysis of Pure Zygotic Genes defined by Baia-Amaral *et al.*, 2024^6^ upon the depletion of *bckdk* mRNA (Bckdk KD) or *phf10* mRNA (Phf10 KD) in zebrafish embryos (see methods for details). **E)** Scatter plot representing the fold change in mRNA level between embryos coinjected with RfxCas13d protein and gRNAs targeting *bckdk* mRNA (y axis) and embryos co-injected with RfxCas13d protein and a gRNA targeting *phf10* mRNA (x axis). Pure Zygotic Genes mRNAs (PZG) determined from Baia-Amaral *et al.*, 2024^6^ data are represented in blue. Dashed lines indicated 1.5-fold in mRNA levels. **F)** Stacked barplots showing the percentage of phenotypes of zebrafish embryos under indicated conditions. *phf10* mRNA isoforms were co-injected (10 pg/embryo) together with RfxCas13d protein (3 ng/embryo) and a mix of 2 gRNAs (1000 pg/embryo) targeting *bckdk* mRNA. *phf10* mRNA isoforms are represented as; T: Threonine, WT version, D: Aspatic acid, mimic the phosphorylation state mediated by Bckdk, A: Alanine, mimic a constitutively non-phosphorylated version. The results are shown as the averages ± standard error of the mean of each developmental stage from two independent experiments. The phenotype selection criteria were the same as described in Fig. 1C. Number of embryos evaluated (n) for each condition is shown (ns = non-significant, **p<0.01, ***p<0.001. ****p<0.0001, χ2-test).

One of the proteins with the lowest level of phosphorylation upon *bckdk* mRNA KD was Phf10/Baf45a (Fig. 5A, red dot). Phf10 is a specific subunit of the Polybromo-associated BAF, pBAF, chromatin remodeling complex that belongs to the SWI/SNF family^36–38^ (Supp. Fig. 7A). Phf10 was less phosphorylated in a threonine (T16) in the *bckdk* mRNA KD embryos compared to the control (Fig. 5A, Supp. Table 2). To address the role of Phf10 during MZT, we targeted its maternal mRNA using CRISPR-RfxCas13d. Interestingly, the embryos co-injected with Cas13d and gRNA targeting *phf10* maternal mRNA displayed similar epiboly defects comparable to the *bckdk* mRNA depletion (Fig. 5B and Supp. Fig. 7D). RNA-seq and qRT-PCR data showed i) a strong degradation of *phf10* mRNA in CRISPR-RfxCas13d injected embryos compared to controls (Cas13d injected only, Fig. 5C and Supp. Fig. 7E), ii) a general downregulation of PZG (Fig. 5C and D) and iii) a positive correlation between PZG transcriptome in *bckdk* and *phf10* mRNA depletions (Fig. 5E, R= 0.3, *p*-value= 1.5e-06, Pearson correlation).

To define whether the phosphorylated state of Phf10 is one of the major driving forces of *bckdk* mRNA KD phenotype, we first defined the lowest dose of ectopic *phf10* mRNAs encoding for either WT or different mutants of Phf10 (T16A and T16D) tagged with an HA that did not trigger a developmental delay (Supp. Fig. 7F and G). Strikingly, the Bckdk depletion phenotype was rescued by the expression of a mutant of *phf10* mRNA containing an aspartic acid in position (T16D) mimicking the Phf10 phosphorylation by Bckdk in WT conditions (Fig. 5F). However, a mutant with an alanine in position T16 (non-phosphorylatable, T16A) was not able to rescue the *bckdk* mRNA KD phenotype (Fig. 5F). Interestingly, the overexpression of Ph10 WT version in the *bckdk* mRNA KD background exacerbated the epiboly defects (Fig. 5F).

Altogether, our data strongly suggest that i) Phf10 is an important factor during MZT contributing to the transcription of the genome during early zebrafish embryogenesis, and ii) a reduced T16 phosphorylation of Phf10 affects its function and can be partially responsible of the *bckdk* mRNA KD developmental phenotype.

## Discussion

While CRISPR-Cas DNA-targeting screenings have been widely used in zebrafish^63–65^ a comprehensive and straight-forward approach to elucidate the role of the maternally provided RNA during early development is required. Following our initial evidence that CRISPR-RfxCas13d efficiently depletes maternal RNA in zebrafish and other vertebrate embryos^27^, we continued to refine and employ this system^28^. Here, we introduce the application of our optimized CRISPR-RfxCas13d system for maternal-RNA-targeted screenings, aiming to uncover new genes regulating early vertebrate development. First, this work demonstrates an increased efficiency and robustness in mRNA level reduction (Fig. 1D, depletion median: 92%, lowest decreased: 58%) using an enhanced gRNA design^28,66^ and purified RfxCas13d protein, which exceeds our original reported efficacy^27^. This suggests that, in general, the lack of phenotype of the screening candidates is not likely due to a low KD efficiency (Fig. 1C and D). Furthermore, our findings indicate that our screening method can identify novel regulators of early embryogenesis. Indeed, we uncovered 8 mRNAs out of 49 screened, whose depletion induced developmental deficiencies (Fig 1C and Supp. Fig. 1B-D). Notably, 7 of these genes were associated with epiboly defects, whereas one (MIBP) was linked to developmental abnormalities later in embryogenesis (Fig. 1B-D). We speculate that chemically modified and more stable gRNAs^67,68^ may help sustain the knockdown and reveal loss-of-function phenotypes later during development. Additionally, we did not observe collateral activity in our knockdowns (Supp. Fig. 3A and B, and Supp. Fig. 4G) and, in fact, the *bckdk* mRNA KD phenotype could be complemented by injecting a non-targetable version of its mRNA (Fig. 2A and B). The absence of collateral activity in zebrafish could be due to our transient approach versus the constitutive expression of CRISPR-RfxCas13d components employed in mammalian cells where RNA-targeting is accumulated over the time^51–53^. Besides, the collateral activity depends on the amount of target, and we cannot rule out that our optimized conditions could induce toxicity when depleting more abundant RNAs than those targeted in this study (from 45 to 768 TPM, median between 0-4 hpf: 164 TPM^40^).

Second, gene expression regulation during the MZT occurs at multiple levels, including chromatin remodeling and modification, transcription, mRNA stability, translation, etc^4,7–15,39^.However, the significance of post-translational regulation and, particularly, protein phosphorylation state in this context remains understudied. Our research reveals a subset of kinases and a phosphatase whose maternal mRNA KD leads to epiboly defects. While our investigations primarily concentrated on Bckdk for its role in ZGA and MZT, we also identified Mknk2a, a kinase potentially regulating ZGA, alongside other kinases potentially implicated in various early developmental processes beyond ZGA. Notably, our screening identified three kinases linked to calcium signaling, with Cab39l and Calm2a exhibiting similar transcriptomic patterns upon KD, hinting at their interconnected molecular functions (Supp. Fig. 1F). Given the established links between calcium signaling and critical developmental milestones such as epiboly, egg activation, fertilization, and cell cycle regulation^42,69^, exploring the roles of these calcium-dependent kinases could enhance our comprehension of early embryogenesis regulation. Consequently, our findings illuminate the crucial role of protein phosphorylation in governing the initial stages of embryogenesis.

Third, through the first phospho-proteome performed at ZGA, we demonstrated that the phosphorylation state of Bckdha, the main target of Bckdk in the mitochondria, was unaltered (Fig. 5A, Supp. Table 2) in Bckdk depleted embryos and consistently BCAA levels were similar to the control condition ^30,47^(Fig. 2 C and D). Besides, functional mitochondria are maternally provided without mitochondrial renovation during MZT^70,71^ and no significant phosphorylation decrease in mitochondrial proteins is detected upon Bckdk depletion. Altogether these data strongly suggest that *bckdk* mRNA KD did not affect the known Bckdk mitochondrial activity^30,47^. Notably, non-mitochondrial Bckdk has a known role in cancer promotion and cell proliferation and migration^31,33,72^ that, in turn, may be related to stem cell and pluripotency-like states^73,74^. Thus, our approach allowed us to uncover a novel and specific role for Bckdk during early zebrafish embryogenesis, since a maternal mutant of this kinase, in the hypothetical case of being viable and fertile^55^, would have also affected its mitochondrial function.

Fourth, for the first time, we have employed SLAM-seq in conjunction with CRISPR-RfxCas13d-mediated KD in zebrafish embryos. This innovative approach has enabled us to comprehensively investigate the role of Bckdk during early embryogenesis. Our findings demonstrate that *bckdk* mRNA KD diminishes the nascent accumulation of PZG and a large set of maternal-zygotic genes that are normally transcribed at 4 hpf (Fig. 3B). Interestingly, we also identified another set of genes that showed an increase in their transcription and a smaller group which experienced an earlier activation. Remarkably, the vast majority of these last genes are usually transcribed during the MZT, indicating that, alongside ZGA, *bckdk* mRNA KD disrupts the transcription dynamics of early expressed genes without affecting others later activated. Up-regulated genes were involved in translation and protein biogenesis suggesting a possible attempt at gene expression compensation. Furthermore, we noted a modest yet detectable alteration in chromatin accessibility, which, did not exhibit a clear bias towards any specific transcription factor binding sites influenced by Bckdk depletion (Fig. 3D). These results suggest that the impact of *bckdk* mRNA KD could extend to general transcription factors or chromatin modifiers, rather than being confined to specific pioneer regulators where changes in the genome are more prominent upon their loss-of-function^8–10^. Indeed, *bckdk* mRNA depletion reduced H3K27ac, a chromatin modification required for ZGA^5,8,35^ (Fig. 3F). CREBBPB among the known proteins potentially involved in H3K27 acetylation in zebrafish^5^ was the only identified in our proteome or phospho-proteome at 4 hpf without significant changes (Supp. Table 2). Thus, we cannot exclude that the activity, level, or phosphorylation status of one or more of these proteins (EP300A, EP300B, CREBBPA and CREBBP) might be affected by the absence of Bckdk. Instead, Phf10/Baf45a, a unique component of the pBAF chromatin remodeling complex, which is part of the broader SWI/SNF family^36–38^ and necessary to maintain the proliferation state of different cell types^75,76^, was identified as a key potential target of Bckdk. Upon Bckdk depletion, Phf10 was less phosphorylated at T16, a residue whose phosphorylation is conserved in human^77^. CRISPR-Cas13d-mediated *phf10* mRNA KD replicated both the developmental phenotype and PZG down-regulation observed with *bckdk* mRNA KD. Remarkably, a phosphomimetic version of Phf10 mutated at T16 rescued the epiboly defects found in embryos depleted on Bckdk. This suggests that one of the main (direct or indirect) Bckdk targets regulating MZT is Phf10 and that Bckdk modulates pBAF activity during early development. Although pBAF is rather a reader of H3K27ac than a complex involved in the deposit of this epigenetic mark, its alteration can lead to a reduced transcription and a lower histone acetylation^78–80^. Interestingly, Phf10 phosphorylation has been reported to affect its stability in mammalian cells^81,82^. However, we did not detect Phf10 in our proteome likely due to the differential level of resolution of proteomic and phospho-proteomic approaches. Notably, the overexpression of Phf10 causes epiboly delay independently of the T16 phosphorylation (Supp. Fig. 7F) status indicating that a correct level of this protein is important to progress through development. In addition to Smarca2 and Smarca4a, which are well-known to be phosphorylated during mitosis to remove them from chromatin we also detected phosphorylation of other SWI/SNF complex members including Arid1a, Arid1b, Arid2, Pbrm1, and Smarcb1, at 4 hpf ^38,83^(Supp. Table 2), suggesting protein phosphorylation could be an important post-translational regulator of the activity of this complex during MZT. Furthermore, our phospho-proteomic analysis detected phosphorylated proteins related to MZT such as Nanog, Brd3a, Brd3b, Brd4 and H1m^5,9,10,27,29,84^. Together, these data suggest that, beyond the regulation of Phf10 mediated by Bckdk, the phosphorylation status of these chromatin and transcription factors may play a crucial role in mediating their function during the ZGA and MZT. Moreover, we identified other proteins exhibiting altered phosphorylation patterns after *bckdk* mRNA KD which may also contribute to disruptions in ZGA. Among these, Znf148 (Supp. Fig. 7A) with a predicted transcription factor activity and Lbh, a well-known transcriptional cofactor^85^

Another interesting candidate is Ago3a, which displayed a decreased phosphorylation of a tyrosine residue in the *bckdk* mRNA KD (Fig. 5A and Supp. Fig. 7A). Other phosphorylated residues in different Ago proteins have been associated with Argonaute–mRNA interactions^86,87^. The temporal reduction of miR-430 processing observed in Bckdk depleted embryos during MZT triggers an increased stability of a subgroup of targets of this miRNA (Fig. 4A). However, Ago3a reduced phosphorylation might also be related with a potential decrease in the regulatory efficacy of miR-430. Intriguingly, we could not find any alteration in the phosphorylation state of Dicer1 or protein levels of Pasha, known proteins involved in miRNA processing detected in our phospho-proteome and proteome, respectively (Supp. Table 2). However, we cannot exclude the possibility that the resolution of these approaches did not allow us to detect changes in other proteins that could be responsible of the delayed biogenesis of miR-430. Further experiments would need to be done to deeply analyze the relation between miRNA processing and protein phosphorylation mediated by Bckdk.

In conclusion, we have been able to reveal that i) CRISPR-RfxCas13d is a powerful tool to perform maternal mRNA KD screenings (49 genes), ii) Bckdk regulates ZGA and MZT not only in zebrafish but also in other teleosts such as medaka and iii) this regulation is, at least, mediated through a reduced biogenesis and activity of miR-430 and a posttranslational phosphorylation of Phf10 whose loss-of-function negatively affects the activation of the zebrafish genome. The influence of Bckdk on the two main events of MZT, zygotic genome activation and maternal mRNA clearance emphasizes the critical role of post-translational phosphorylation as a regulatory mechanism during embryogenesis and underscores the importance of further exploring the functions of kinases and phosphatases controlling early development.

## Methods

### Zebrafish Maintenance and Embryo Production

All experiments conducted using zebrafish at CABD adhere to national and European Community standards regarding the ethical treatment of animals in research and have been granted approval by the Ethical committees from the University Pablo de Olavide, CSIC and the Andalusian Government. Zebrafish wild type strains AB/Tübingen (AB/Tu) were maintained and bred under standard conditions^88^. Wild-type zebrafish embryos were obtained through natural mating of AB/Tu zebrafish of mixed ages (5–18 months). Selection of mating pairs was random from a pool of 10 males and 10 females. Zebrafish embryos were staged in hpf as described^89^. Zebrafish experiments at Stowers Institute were done according to the IACUC approved guidelines. Zebrafish embryos for microinjections were coming from random parents (AB, TF and TLF, 6–25 months old) mating from 4 independent strains of a colony of 500 fish. The embryos were pooled from random 24 males and 24 females for each set of experiments.

### Medaka Maintenance and Embryo Production

All experiments involving medaka adhered to national and European Community standards for animal use in research and received approval from the Ethical Committees of the University Pablo de Olavide, CSIC, and the Andalusian Government. The wild type medaka strain (iCab) was kept and bred under standard conditions. Fifteen to twenty pairs of four-month-old males and females were randomly selected and crossed to produce embryos for the experiments described here. Embryos were injected at the single-cell stage following standard methods. Their progression through developmental stages was determined in hours post-fertilization (hpf) using the described procedures^90^.

### Selection of candidates for maternal screening

Maternal screening candidates were selected according to their levels of maternally provided RNA, translation levels, and functional annotation. mRNAs highly maternally provided (> 50 TPM at 0 hpf in rib0, Paper Ariel 2021) and high translated at 2 hpf (>130 rpkm in ribosome profiling data^5,39^) were soft clustered by Mfuzz Software in R (v. 2.62.0)^91^ using ribosome profiling data at 0, 2, 5 and 12 hpf^5,39^. mRNA following a dynamic of highly translation between 0-5 hpf and then a decrease after this timepoint were chosen. Among then, mRNA encoding kinases or phosphatases based on gene description by ZFIN database information^92^ were selected for the maternal screening.

### Guide RNA design RfxCas13d protein and mRNA production

All Guide RNAs (gRNAs) and Cas13d protein (RfxCas13d) employed in this study were design according to previous detailed protocol for zebrafish embryos^28^. 3 gRNAs targeting the CDS were design per each mRNA. All designed gRNAs for maternal screening were chemically synthesized by Synthego (Synthego Corp., CA, USA). gRNAs for phf10 mRNA knockdown were generated by fill-in-PCR with a universal primer and 3 specific primers (Supp. Table 1) with the Q5 High-Fidelity DNA Polymerase. Purify PCR product using FavorPrep Gel/PCR Purification Kit were used as template for *in vitro* gRNA production using AmpliScribe T7 Flash Transcription Kit (Epicentre,Lucigen # ASF3257). Then gRNAs were precipitated according to previous detailed protocol and resuspended in nuclease free water^28^. gRNAs were quantified using Qubit RNA BR (Broad-Range) assay kit (ThermoFisher, #Q10210).

Constructions for *bckdk* and *phf10* mRNA rescue and overexpression experiments were designed by cloning CDS sequence with specific primers (Supp. Table 1) using cDNA from 4 hpf or purchasing the CDS sequence from Integrated DNA Technologies (https://eu.idtdna.com/), respectively. Different phosphorylation state isoforms for *phf10* mRNA were generated by site-directed mutagenesis using QuikChange Multi Site-Directed Mutagenesis kit (#200514, Agilent), following manufacturer’s instructions and specific primers (Supp. Table 1). Plasmid constructions (Supp. Table 1) were digested with *XbaI* for 2h at 37°C to linearize the DNA and then used for in vitro transcription using the mMESSAGE mMACHINE™ T3 (#AM1348, ThermoFisher Scientific) following manufacturer’s protocols. mRNA products were purified using the RNeasy Mini Kit (#74104, Qiagen) and quantified using Nanodrop.

### Zebrafish embryo injection and image acquisition

One-cell stage zebrafish embryos were injected with 1 nL containing 3 μg/μL of purified RfxCas13d protein and 1000 ng/μL of gRNAs (individual or a mix of 2-3 gRNAs, see figure legends for details in each experiment). Between 10 to 50 pg per embryo of ectopic *gfp* and *bckdk* mRNA were injected for rescue experiment. Between 10 to 50 pg of each version of *phf10* mRNA were injected for overexpression or rescue experiments. miR430 duplexes (Supp. Table 1) were purchased from IDT (https://eu.idtdna.com/) and resuspended in RNAse-free water. 1 nL of 2.5 µM miR-430 duplex solution (equimolar mix of 3 duplexes miR-430a, miR-430b and miR-430c) were injected at one-cell-stage, using single-use aliquots. Zebrafish embryos phenotypes were quantified at 2, 4, 6 and 24 hpf. Zebrafish embryo phenotypes pictures were performed using a Nikon DS-F13 digital camera and images processed with NIS-Elements D 4.60.00 software. GFP fluorescence signal were quantified using Fiji (Image J) software, 15 embryos in 3 images (5 embryos/image) per condition were employed.

### Medaka embryo injection and image acquisition

One-cell stage medaka embryos were co-injected with 2–3 nL of a solution containing 3 μg/μL of purified RfxCas13d protein and 1000 ng/μL gRNA. Medaka embryos phenotypes were quantified at 24 and 48 hpf. Medaka embryo phenotypes pictures were performed using a Nikon DS-F13 digital camera and images processed with NIS-Elements D 4.60.00 software.

### RT-qPCR

Total RNA to analyse mRNA levels by RT-qPCR were extracted from 10 embryos per biological replicate. Embryos were collected at the specific timepoint and snap-frozen in liquid nitrogen. Total RNA was extracted using standard TRIzol protocol as described in the manufacturer’s instructions (ThermoFisher Scientific). Then, cDNA was generated using 1000 ng of total purified RNA following iScript cDNA synthesis kit (1708890, Bio-Rad) manufacturer’s protocol. 2 µl of a 1:5 cDNA dilution was used together with forward and reverse primers per each mRNA (2 µM; Supp. Table 1) and 5 µl of SYBR® Premix-Ex-Taq (Tli RNase H Plus, Takara) in a 10 µl reaction. PCR cycling profile consisted in a denaturing step at 95 °C for 30 s and 40 cycles at 95 °C for 10 s and 60 °C for 30 s. *taf15* mRNA was used as control for sample normalization.

### Primary-miR-430 and mature-miR-430 quantification

10 embryos per biological replicate were collected at 4.3 and 6 hpf to analyse by RT-qPCR primary miR-430 and mature miR430 levels. Embryos were snap-frozen in liquid nitrogen at indicated timepoint. Total RNA and RNA enriched in small RNA species were extracted using mirVana™ miRNA Isolation Kit (#AM1561, ThermoFisher Scientific) following manufacturer’s instructions. cDNA for measuring primary-miR430 was generated using 200 ng of total purified RNA following iScript cDNA synthesis kit (#1708890, Bio-Rad) manufacturer’s protocol. Then, 2 µl of cDNA was used together with specific forward and reverse primers for primary miR-430^93^ (2 µM; Supp. Table 1) and 5 µl of SYBR® Premix-Ex-Taq (Tli RNase H Plus, Takara) in a 10 µl reaction. *taf15* mRNA was used as control for sample normalization.

100 ng of enriched small RNA fraction was used for generated cDNA following iScript Select cDNA Synthesis Kit (#708896, Bio-Rad) following manufacture’s protocol and the specific primer 5′-GCAGGTCCAGTTTTTTTTTTTTTTTCTACCCC-3′ (Supp. Table 1). 2 µl of cDNA was used together with specific primers for each mature miR-430 (a, b and c) and the previous primer (2 µM; Supp. Table 1), and 5 µl of SYBR® Premix-Ex-Taq (Tli RNase H Plus, Takara) in a 10 µl reaction. ncRNA u4atac^94^ was used for sample normalization in mature microRNA RT-qPCR.

### RNA-seq Libraries and Analysis

20 zebrafish embryos per biological replicate were collected at 4 or 6 hpf and snap-frozen in 300 µl of DNA/RNA Shield (#R1100-50, Zymo Research). Total RNA then was extracted using Direct-zol RNA Miniprep Kit (#R2050, Zymo Research) following manufacturer’s instructions. 10 medaka embryos per biological replicate were collected and dechorionated as described in Porazinski et al., 2010 at 6.5 or 8 hpf. Total RNA was extracted using standard TRIzol protocol as described in the manufacturer’s instructions (ThermoFisher Scientific). Total RNA from zebrafish and medaka embryos was quantified using the Qubit fluorometric quantification (#Q10210, Thermo).

For RNA-seq libraries for maternal screening associated to Figure 1 and Supp. Fig 2, cDNA was generated from 1.25 ng of high-quality total RNA (except for Cas13d control and *bckdk* mRNA KD where cDNA was generated from 2.5 ng of high-quality total RNA), as assessed using the Bioanalyzer (Agilent), according to manufacturer’s directions for the SMART-seq v4 Ultra Low Input RNA Kit (Takara, 634891) at a 1/8th reaction volume and using the Mantis (Formulatrix) nanoliter liquid-handling instrument to pipette the reagents for cDNA synthesis (except for Cas13d control and *bckdk* mRNA KD samples where 1/4th of the reaction volume were used). Libraries were generated manually (except for cab39l and calm2a mRNA KD samples where libraries were generated with the Mosquito HV Genomics (SPT Labtech) nanoliter liquid-handling instrument), using the Nextera XT DNA Library Preparation Kit (Illumina, FC-131-1096) at 1/8th reaction volumes (except for Cas13d control and *bckdk* mRNA KD samples where 1/4th of the reaction volume were used) paired with IDT for Illumina DNA/RNA UD Indexes Set A (Illumina, 20027213), and purified using the Ampure XP bead-based reagent (Beckman Coulter, Cat. No. A63882). Resulting short fragment libraries were checked for quality and quantity using the Bioanalyzer and Qubit Fluorometer (ThermoFisher). Equal molar libraries were pooled, quantified, and sequenced on a High-Output flow cell of an Illumina NextSeq 500 instrument using NextSeq Control Software 2.2.0.4 with the following read length: 70 bp Read1, 10 bp i7 Index and 10 bp i5 Index. Following sequencing, Illumina Primary Analysis version NextSeq RTA 2.4.11 and Secondary Analysis version bcl2fastq2 (v. 2.20) were run to demultiplex reads for all libraries and generate FASTQ files. Exception: libraries for Ppp4r2a KD employed in the maternal screening in Figure 1 were performed similarly to libraries for medaka samples and Phf10 KD associated to Figure 2 and Figure 5 (see below).

RNA-seq libraries for medaka samples and *phf10* mRNA KD associated to Figure 2 and Figure 5, respectively were generated from 100 ng (or ≤100 ng) of high-quality total RNA, as assessed using the Bioanalyzer (Agilent), were made according to the manufacturer’s directions using a 25-fold (or 100-fold) dilution of the universal adaptor and 13 cycles (or 16 cycles) of PCR per the respective masses with the NEBNext Ultra II Directional RNA Library Prep Kit for Illumina (NEB, Cat. No. E7760L), the NEBNext Poly(A) mRNA Magnetic Isolation Module (NEB, Cat. No. E7490L), and the NEBNext Multiplex Oligos for Illumina (96 Unique Dual Index Primer Pairs) (NEB, Cat. No. E6440S) and purified using the SPRIselect bead-based reagent (Beckman Coulter, Cat. No. B23318). Resulting short fragment libraries were checked for quality and quantity using the Bioanalyzer and Qubit Flex Fluorometer (Life Technologies). Equal molar libraries were pooled, quantified, and converted to process on the Singular Genomics G4 with the SG Library Compatibility Kit, following the “Adapting Libraries for the G4 – Retaining Original Indices” protocol. The converted pool was sequenced on an F3 flow cell (Cat. No. 700125) on the G4 instrument with the PP1 and PP2 custom index primers included in the SG Library Compatibility Kit (Cat. No. 700141), using Instrument Control Software 23.08.1-1 with the following read length: 8 bp Index1, 100 bp Read1, and 8 bp Index2. Following sequencing, sgdemux (v. 1.2.0) was run to demultiplex reads for all libraries and generate FASTQ files.

All raw reads were demultiplexed into Fastq format allowing up to one mismatch using Illumina bcl2fastq2 v2.18. Reads were aligned to UCSC genome danRer11 or to UCSC genome oryLat2 with STAR aligner (v. 2.7.3a), using Ensembl 102 gene models. TPM values were generated using RSEM (v. 1.3.0). Fold change for each gene was calculated using Deseq2 package (v. 1.42.0) after filtering genes with a count of 20 reads in all control libraries. Differential expression genes were selected setting a corrected p-value < 0.05 and a fold change > 3. Common downregulated genes were calculated using UpSet intersection visualization the R package UpSetR^95^(v. 1.4.0)

RNA-seq libraries for samples at 6 hpf in zebrafish embryos associated to Figure 4 were generated from 100 ng of high-quality total RNA, as assessed using the Bioanalyzer (Agilent). Libraries were made according to the manufacturer’s directions for the TruSeq Stranded mRNA Library Prep Kit (Illumina, Cat. No. 20020594), and TruSeq RNA Single Indexes Sets A and B (Illumina Cat. No. 20020492 and 20020493) and purified using the Ampure XP bead-based reagent (Beckman Coulter, Cat. No. A63882). Resulting short fragment libraries were checked for quality and quantity using the Bioanalyzer (Agilent) and Qubit Fluorometer (Life Technologies). Equal molar libraries were pooled, quantified, and sequenced on a High-Output flow cell of an Illumina NextSeq 500 instrument using NextSeq Control Software (v. 4.0.1) with the following read length: 76 bp Read1, 6 bp i7 Index. Following sequencing, Illumina Primary Analysis version NextSeq RTA 2.11.3.0 and bcl-convert-3.10.5 were run to demultiplex reads for all libraries and generate FASTQ files.

### Slam-seq Libraries and Analysis

s4-UTP (SLAMseq Kinetics Kit – Anabolic Kinetics Module; #061, Lexogen) were injected at 25 50 and 75 pmol/embryo in one-cell-stage embryos under red light and kept in the dark until desired timepoint. Due to toxicity effect of s4-UTP in combination of RfxCas13d protein, 25mM aliquots were selected as final concentration for Slam-Seq experiment. This reduced levels of s4-UTP could potentially compromise the coverage of label reads compared to optimized conditions in zebrafish embryos^6,18,59^. 25 embryos were collected at 4 hpf and snap-frozen in 500 µl of Trizol in tubes protected from light. Total RNA was extracted and then alkylated with Iodoacteamide under dark conditions using SLAMseq Kinetics Kit – Anabolic Kinetics Module (#061, Lexogen) following manufacturer’s instructions.

3′-end mRNA sequencing libraries were generated, according to the manufacturer’s instructions, from 200 ng of alkylated total RNA (except no-s4U Cas13d-alone sample, which was 87ng), using the QuantSeq 3′ mRNA-seq Library Prep Kit for Illumina UDI Bundle (FWD, Lexo gen GmbH, cat. no. 144.96). ERCC-92 RNA spike-ins were added at equal final molarity to all samples. For PCR amplification, 14 cycles were performed with the Lexogen Unique Dual Index (UDI) 12-nt Index Set B1. The resulting libraries were checked for quality and quantity using the Qubit Fluorometer (Life Technologies) and the Bioanalyzer (Agilent). Libraries were pooled, re-quantified, and sequenced as 100-bp single reads on the NextSeq 2000 (Illumina). Following sequencing, Illumina Primary Analysis RTA and bcl2fastq2 were run to demultiplex reads for all libraries and generate FASTQ files. Adapters were cut from FASTQ files using TRIMGALORE with the following parameters: -a AGATCGGAAGAGCACACGTCTGAACTCCAGTCAC; --length 30. All trimmed reads were mapped to danRer11 genome using default parameters of slamdunk map module (v. 0.4.3), then filtered with filter module. Zebrafish ENS102 annotation for protein coding genes 3′UTRs and non-coding genes coordinates were used as reference to get reads per 3′UTR or non-coding gene. To estimate background T>C due to SNPs, a VCF file was obtained from Cas13d alone (no-S4U) and Cas13+gRNAs (no-s4U) with slamdunk snp module with following parameters: -f 0.2. Only then, total Reads and labelled reads (containing at least 2 T>C conversion events) were obtained with slamdunk count module using the VCF file created in previous steps (unlabelled reads were estimated by subtracting labelled from total read counts).

Read counts per million (CPM) were obtained using Deseq2 (v. 1.42.0). Fold change for each gene and its associated *p*-value was calculated using Deseq2 package (v. 1.42.0) after filtering genes with a count of 20 reads in all control libraries for unlabelled and labelled data. One replicate from Bckdk KD and 3 from Cas13d control were discarded from the analysis due to their extremely low sequence depth and spike-ins over-representation.

### Gene Set Enrichment Analysis and Gene Ontology Analysis

Differential expression analysis from RNA-seq or Slam-seq data generated were ranked according to its fold change compared to control conditions and used as input for Gene Set Enrichment Analysis^96,97^ (GSEA) resulting in Normalized Enrichment Score with its adjusted *p*-value. GSEA analyses were performed using the R Package clusterProfiler (v. 4.10.0). Conditions with a GSEA Normalized Enrichment score <-2 or >2 and a p-adjusted < 0.01 were considered as significant.

Enrichment of Gene Ontology Biological Process terms was calculated using ShinyGO^98^ terms with a false discovery rate (FDR), corrected p-value < 0.05 and with more than 20 genes represented are considered as enriched.

### ATAC-seq Libraries and Analysis

ATAC-seq assays were conducted following standard protocols^99,100^, with slight adjustments. A total of 80 embryos per experimental condition were collected at 4 hpf and dissolved in Ginzburg Fish Ringer buffer without calcium (55 mM NaCl, 1.8 mM KCl, 1.15 mM NaHCO3) by gentle pipetting and shaking at 1100 rpm for 5 minutes. The samples were then centrifuged for 5 minutes at 500g at 4°C. Following removal of the supernatant, the embryos were washed with cold PBS and subsequently resuspended in 50 μl of Lysis Buffer (10 mM Tris–HCl pH 7.4, 10 mM NaCl, 3 mM MgCl2, 0.1% NP-40) by gentle pipetting. The equivalent volume of 70,000 cells was centrifuged for 10 minutes at 500g at 4°C, after which it was resuspended in 50 μl of Transposition Reaction mixture, comprising 1.25 μl of Tn5 enzyme and TAGmentation Buffer (10 mM Tris–HCl pH 8.0, 5 mM MgCl2, 10% w/v dimethylformamide), followed by an incubation period of 30 minutes at 37°C. Subsequent to TAGmentation, DNA was purified using the MinElute PCR Purification Kit (Qiagen) and eluted in 10 μl. Libraries were generated via PCR amplification using NEBNext HighFidelity 2× PCR Master Mix (NEB). The resultant libraries were purified again using the MinElute PCR Purification Kit (Qiagen), multiplexed, and then sequenced on a HiSeq 4000 pair-end lane, yielding approximately 100 million 49-bp pair end reads per sample.

ATAC-seq reads were aligned to the GRCz11 (danRer11) zebrafish genome assembly using Bowtie2 2.3.5 and those pairs separated by more than 2 kb were removed. The Tn5 cutting site was determined as the position -4 (minus strand) or +5 (plus strand) from each read start, and this position was extended 5 bp in both directions. Conversion of SAM alignment files to BAM was performed using Samtools 1.9^101^. Conversion of BAM to BED files, and peak analyses, such as overlaps or merges, were carried out using the Bedtools 2.29.2 suite (51). Conversion of BED to BigWig files was performed using the genomecov tool from Bedtools and the wigToBigWig utility from UCSC^102^

ATAC peaks were called using MACS2 and peaks were pool together to calculate fold change for each peak using Deseq2 package (v1.3.0). Differential accessibility regions were selected setting a corrected p-value < 0.05 and a fold change > 2-fold. Bed files were normalized according to relative number of reads into peaks. Heatmaps and average profiles of ATAC-Seq data were performed using computeMatrix, plotHeatmap and plotProfile tools from the Deeptools 3.5 toolkit. Transcription Factor motifs were calculated using Gimmemotifs (v. 0.18.0) with standard parameters. Differential ATAC peak groups (setting a corrected P value < 0.05 and a fold-change > 1.5 or 3-fold) from danRer 11 were converted to danRer7 using the Liftover tool of the UCSC Genome Browser^103^. These peaks were assigned to genes using the GREAT 3.0.0 tool with the basal plus extension association rule with the following parameters: 5 kb upstream, 1 kb downstream, 1000kb of maximum extension. These genes were associated with Slam-Seq differential labelled data and represented as a violin plot for those conditions with > 10 genes per group.

### Leucine-isoleucine and valine quantification

To analyse aminoacid levels, 50 embryos were collected at 4 hpf and resuspended in 1ml of Deyolking Buffer (55mM NaCl, 1.8mM KCl, 1.25mM NaHCO3). Samples were incubated at 25°C for 5 minutes with orbital shaking at 1100 rpm and then centrifuge at 300G for 30s. Supernatant was removed and samples were washed with 1ml of Wash Buffer (110mM NaCl, 3.5mM KCl, 2.7mM NaHCO3) shaking the tubes at 1100 rpm for 2 min. Samples were centrifuge at 300G for 30 s and the supernatant was resuspended in 350 ul of methanol with 10 µg/ml of leucine used as internal control measure or in 0.1N HCl for valine analysis. Samples were sonicated for 15 min at 4°C to enhance the supernatant and centrifuge again. Finally, samples were centrifuge for 10 min at 16.000G and the supernatant were stored at -80°C until samples processing.

Leucine and isoleucine (Sigma-Aldrich), 13C6-Leucine (Cambridge Isotopes), valine (Sig-ma-Aldrich), L-Valine-1-13C (Sigma-Aldrich) and samples were derivatized by dansyl chloride (Sigma-Aldrich). 3.5 mM amino acids were dissolved in 20 µl of pH 9.5 250 mM sodium bicarbonate buffer, 16 times more dansyl chloride was added and incubated at room temperature for 1 hour. The reaction was quenched by adding 11 µl of 0.5 M heptylamine (Sigma-Aldrich) at room temperature for 15 minutes. 40 µl of each sample was dried by vacuum centrifugation and reconstituted with 20 µl of 250 mM pH 9.5 sodium bicarbonate buffer, then mixed with 2 µl 56mM dansyl chloride for 1 hour at room temperature. The reaction was quenched by adding 11 µl of 0.5 M heptylamine for 10 minutes at room temperature. 400 nM of derivatized 13C6-Leucine was spiked into 2, 10, 25, 50, 75, 100 nM of derivatized Leucine respectively, as well as each derivatized sample before LC/MS analysis. Amino acid samples were analyzed on a Q-Exactive Plus Mass Spectrometer (Thermo Scientific) equipped with a Nanospray Flex Ion Source and coupled to a Dionex UltiMate 3000 RSCLnano System. A 75 µm i.d. analytical microcapillary column was packed in-house with 100 mm of 1.9 μm ReproSil-Pur C18-AQ resin (Dr. Masch). AgileSLEEVE (Analytical Sales & Products) was used to maintain column temperature at 50°C. The UPLC solutions were 0.1% formic acid in water for buffer A (pH 2.6) and 0.1% formic acid in acetonitrile for buffer B. The chromatography gradient as a 5-min loading time at 20%B, 23 min from 20% to 35%; 2 min to reach 100% B; and 10 min washing at 100% B. The nano pump flow rate was set to 0.5 ul/min. The Q-Exactive was set up to run a Parallel Reaction Monitoring (PRM) method with inclusion list as 371.1736 m/z and 365.1535 m/z with HCD at 40%. Each standard curve concentration points and each sample were analyzed in triplicates. The peak area ratio between leucine and 13C6-leucin and the peak ration between valine and L-valine-1-13C was used to quantify the leucine and valine amount in each sample.

### Protein sample Preparation and Western Blot

20-25 embryos were collected at 4 or 6 hpf and washed with 200 µl of Deyolking Buffer (55mM NaCl, 1.8mM KCl, 1.25mM NaHCO3). Then, embryos were lysated adding other 200 µl of Deyolking Buffer and by pipetting up and down. Samples were incubated at 25°C for 5 minutes with orbital shaking and centrifuge at 300G for 30’’. Supernatant were removed and the pellet was washed by adding 300 µl of Wash Buffer (110mM NaCl, 3.5mM KCl, 10mM TRIS-HCl pH= 7.4). Samples were centrifuged at 300G for 30 s and the supernatant was resuspended in 10 µl of SDS-PAGE buffer (160 mM Tris-HCl pH 8, 20% Glycerol, 2% SDS, 0.1% bromophenol blue, 200 mM DTT). SDS-PAGE electrophoresis was conducted using 10% or 12% TGX Stain-FreeTM Fast CastTM Acrylamide Solutions (Bio-Rad). Subsequently, the protein gels were activated with a Chemidoc MP (Bio-Rad) and transferred to Nitrocellulose membranes via the Trans-Blot Turbo Transfer System (Bio-Rad). Following this, the membranes were blocked at room temperature for 1 hour using Blocking Solution (5% fat-free milk in 50 mM Tris-Cl, pH 7.5, 150 mM NaCl (TBS) with 1% Tween20).

The primary antibody Anti-H3K27Ac (#ab177178, Abcan) was diluted 1:5000, Anti-HA (#11867423001, Roche) was diluted 1:1000 and Anti-H3 (#sc-517576, Santa Cruz Biotechnology) was diluted 1:500, while the secondary antibody, anti-mouse HRP-labelled (#A5278, Sigma-Aldrich), was diluted 1:5000 and anti-rabbit HRP-labelled (#ab6721, Abcam) was diluted 1:2000, all in Blocking Solution.

The membrane was then incubated with the primary antibody overnight at 4 °C. Post-primary antibody incubation, the membrane underwent three brief washes with TBS with 1% Tween 20 (TTBS) for 15 minutes each, followed by incubation with the secondary antibody for 60 minutes at room temperature. Washes were performed similarly to those with the primary antibody. Detection was accomplished using ClarityTM Western ECL Substrate (Bio-Rad), and images were captured with a ChemiDoc MP (Bio-Rad). H3K27ac and H3 intensity were calculated using ImageLab software and normalized to total Stain-Free total lane volumes^104^.

### Protein extraction for proteomic analysis

100 embryos collected at 4 hpf were resuspended in 1 ml of Deyolking Buffer (55mM NaCl, 1.8mM KCl, 1.25mM NaHCO3) by shaking the tubes at 1100 rpm for 5 min. Samples were centrifuged at 300G for 30 s. Supernatant were removed and pellet washed with 1 ml of Wash Buffer (110mM NaCl, 3.5mM KCl, 2.7mM NaHCO3) and shaken for 2 min at 1100 rpm. After centrifugation at 300 G for 30 s, samples were resuspended in homogenize buffer (50 mM Tris (pH 7.5), 1 mM DTT, Phosphatase|Protease inhibitor EDTA free). Samples were briefly sonicated (twice for 10 s) and centrifuged at 16,900G for 10 min at 4°C. The supernatant was kept it in ice and protein concentration were measured with QUBIT protein kit (#Q33211, ThermoFisher).

### Digestion for Mass Spectrometry

Samples were reduced by adding tris(2-carboxyethyl) phosphine (TCEP at 1M) to 5 mM final at room temperature for 30 minutes. To carboxymethylate reduced cysteine residues, 5 μl of 2-chloroacetamide (CAM, made fresh at 0.5 M) were added and samples were incubated for 30 minutes protected from light at room temperature. Six volumes of pre-chilled (at -20°C) acetone were added and the precipitation was let to proceed overnight. Samples were next centrifuged at 8000 × g for 10 minutes at 4°C. The tubes were carefully inverted to decant the acetone without disturbing the protein pellets, which were air-dried for 2-3 minutes.

The precipitated proteins were dissolved in 100 μl of 50 mM TEAB, mass spectrometry grade trypsin (Promega Gold) was added at 1:40 w/w and the digestion were let to proceed at 37°C overnight. The digested protein samples were centrifuged at 16000 x g for 30 minutes and transferred to new tubes. A fluorometric peptide assay (Pierce) was performed on 10 μl of each digested sample according to manufacturer’s instructions. Peptide amounts ranged from 30 to 120 μg.

### Tandem Mass Tag (TMT) Labeling

Immediately before use, the vials containing frozen TMTpro 16plex reagents (thermo Scientific) were let to warm up at room temperature. To each of the 0.5mg vials, 20 µl of anhydrous acetonitrile were added. The TMT reagents were let to dissolve for 5 minutes with occasional vortexing, then the tubes were briefly centrifuged to gather the solution. For each sample, 20 μg were measured out based on the peptide assay, and the volumes were adjusted to 100 µl with 100 mM TEAB, then mixed with the 20 µl in TMT vials. The 4 replicate samples for the Cas13d control condition and Bckdk KD condition, were labeled with TMTpro--129N, -129C, -130C -130C, -131N, - 131C, -132N, and -132C respectively. The labeling reaction was allowed to proceed for 1 hour at room temperature. To test labeling efficiency, 1 µl of samples were analyzed by LC/MSMS over a 2-hour C18 reverse-phase (RP) gradient on an Orbitrap Eclipse Tribrid Mass Spectrometer (Thermo Scientific) with a FAIMS Pro interface, equipped with a Nanospray Flex Ion Source, and coupled to a Dionex UltiMate 3000 RSCLnano System. The TMT labeling levels were >99% for all detected peptides (data not shown). The 12 differentially labeled samples (30 µl each) were combined in a new tube and the resulting volume was reduced using a SpeedVac concentrator (Savant) to less than 10 µl (about 2 hour).

### Phosphopeptide Enrichment

High-SelectTM Fe-NTA Phosphopeptide Enrichment Kit (Thermo Scientific) was used for phosphopeptide enrichment. Lyophilized TMT labeled peptides were suspended in 200 μl of binding buffer and loaded onto equilibrated Fe-NTA spin column. After 30 minutes incubation, the supernatant was collected through centrifugation at 1000 x g. 100 µl of elution buffer were added to the spin column, and the eluate was collected through centrifugation at 1000 x g. Both supernatant and eluate were dried immediately in SpeedVac concentrator to less than 10 µl.

### High pH Reverse Phase Fractionation

The dried supernatant TMT-labeled peptide mixture was resuspended in 300 µl of 0.1% trifluoroacetic acid (TFA). One high pH fractionation cartridge (Pierce, cat. # 84868) was placed on a new 2.0 ml sample tube and 300 µl of the TMT-labeled peptide mixture were loaded onto the column. After centrifuging at 3000 × g for 2 minutes, the eluate was collected as the “flow-through” fraction. The loaded cartridge was placed on a new 2.0 ml sample tube, washed with 300 µl of ddH20, and the eluate collected as “wash” fraction. An additional round of washing was performed using 300 µl of 5% acetonitrile in 0.1% TFA to remove unreacted TMT reagent. A total of 8 HpH RP fractions were collected by sequential elution in new sample tubes using 300 µl of 10%, 12.5%, 15%, 17.5%, 20%, 22.5%, 25%, and 50% acetonitrile in 0.1% TFA. The solvents were evaporated to dryness using vacuum centrifugation. Dried samples were resolubilized in 44 µl of buffer A (5% acetonitrile in 0.1% formic acid, FA) before LC-MS analysis.

### Multiplexed Mass Spectrometry Analysis

TMT-labeled peptides were analyzed on Orbitrap Eclipse Tribrid Mass Spectrometer (Thermo Scientific) with a FAIMS Pro interface, equipped with a Nanospray Flex Ion Source, and coupled to a Dionex UltiMate 3000 RSCLnano System. Peptides (22 µl for each HpH RP fraction) were loaded on an Acclaim PepMap 100 C18 trap cartridge (0.3 mm inner diameter (i.d.), 5 mm length; Thermo Fisher Scientific) with the U3000 loading pump at 2 µL/minute via the autosampler.

A 75 µm i.d. analytical microcapillary column was packed in-house with 250 mm of 1.9 μm ReproSil-Pur C18-AQ resin (Dr. Masch). AgileSLEEVE (Analytical Sales & Products) was used to maintain column temperature at 50°C. The organic solvent solutions were water:acetonitrile:formic acid at 95:5:0.1 (volume ratio) for buffer A (pH 2.6) and 20:80:0.1 (volume ratio) for buffer B. The chromatography gradient was a 30-min column equilibration step in 2% B; a 5-min ramp to reach 10% B; 90 minutes from 10% to 40% B; 10 min to reach 90% B; a 5-min wash at 90% B; 0.1 min to 2% B; followed by a 15-min column re-equilibration step in 2% B. The nano pump flow rate was set to 0.180 μl/min.

The Orbitrap Eclipse was set up to run the TMT-SPS-MS3 method with three FAIMS compensation voltages (CV) at -40V, -55V, and -70V and a cycle time of 1 sec. Briefly, peptides were scanned from 400-1600 m/z in the Orbitrap at 120,000 resolving power before MS2 fragmentation by CID at 35% NCE and detection in the ion trap set to turbo detection. Dynamic exclusion was enabled for 45s. The supernatant from phosphorylation peptide enrichment was analyzed with real time search (RTS) enabled. During LC/MS data acquisition, ddMS2 spectra was searched against a non-redundant Danio rerio sequence database downloaded from NCBI 2021-03 and complemented with common contaminants. Carbamidomethyl (max = 57.0215 Da at C sites) and TMTpro16plex (max = 304.2071 Da at Kn sites) were searched statically, while methionine oxidation (15.9949 Da) was searched as a variable modification. Synchronous precursor scanning (SPS) selected the top 10 MS2 peptides for TMT reporter ion detection in the Orbitrap using HCD fragmentation at 65% NCE at 50,000 resolving power.

### MS/MS Data Processing

The LC/MSn dataset was processed using Proteome Discoverer 2.4 (Thermo Fisher Scientific). MS/MS spectra were searched against a zebrafish protein database (NCBI 2021-03) complemented with common contaminants. SEQUEST-HT implemented through Proteome Discoverer was set up as: precursor ion mass tolerance 10 ppm, fragment mass tolerance 0.3 Dalton, up to two missed cleavage sites, static modification of cysteine (+57.021 Da), and lysine and peptide N-termini with TMT tag (+304.2071 Da) and dynamic oxidation of methionine (+15.995 Da), and phosphorylation of serine, threonine, tyrosine (+79.9663 Da). Results were filtered to a 1% FDR at peptides levels using Percolator through Proteome

### Phosphoproteomic and Proteomic differential analysis

Normalized TMT (Tandem Mass Tag) Intensity readings were used for differential analysis by edgeR (v. 3.19). Protein accession IDs were mapped to Ensembl Genes with biomaRt (v. 2.54.1). Differential phosphopeptides were selected setting a p-value < 0.05 and a fold-change > 2.0. Localization of differential phosphopeptides upon bckdk mRNA knockdown was determined by uniprot^105^.

Mass spectrometry data was normalized by NormalyzerDE54 (v. 3.18). Fold change for each protein was calculated using limma55 (v. 3.54.2) and resulting *p*-values were adjusted with Benjamini-Hochberg method. Differential proteins were selected setting a p-value < 0.05 and a fold-change > 2.0 after filtering proteins present in all control samples.

### Immunofluorescence

Dechorionated embryos at 4 hpf were fixed overnight with 4% PFA at 4 °C in agitation. Then, they were washed twice with PBS at room temperature (RT) and transferred to a 24-well plate and incubated overnight at 4 °C with shaking in 400 µL of primary antibody solution: DAPI 1:1000 (D9542-1g, Sigma-Aldrich), Phalloidin 1:100 (#A12379, Invitrogen), DMSO 5% in PBST (0.8% Tween 20, #). Afterwards, embryos were rinsed twice during 10 min with PBST at RT.

Image acquisition was performed in confocal microscopy Zeiss LSM 880 Airscan with 40x oil objective and coupled to an Andor EMCCD iXOn DU 879 camera. Fluorescence images were then processed with Fiji (ImageJ). For analysing the number of total cells and mitotic cells, 2 different Z-stack were selected from each embryo, using at least 4 embryos per independent experiment.

### Statistical analysis

Statistical analyses were conducted without predefining the sample size. The experiments were carried out without randomization, and investigators were aware of allocation during both experiments and outcome assessment. No data were excluded from the analysis. Number of embryos, replicates and experiments are indicated in figure legends.

For phenotypes derived from embryo microinjections, Xi-square statistical analyses were undertook using rstatix (v. 0.7.2) in R package. Unpaired two-tailed Mann–Whitney test was used to compare Leucine-Isoleucine or Valine levels in control vs Bckdk KD embryos using Prism (GraphPad Software, La Jolla, CA, USA).

For western-blot quantification analysis *p*-value was calculated using one Welch’s t-test with Prism (GraphPad Software, La Jolla, CA, USA). Quantitative RT-qPCR, RIN analysis and GFP intensity p-values were calculated using unpaired t-test or one-way ANOVA test with Prism (GraphPad Software, La Jolla, CA, USA) and Standard Error of the Mean (SEM) was used to show error bars. Grubbs’s test was performed first for outlier’s identification. Unpaired two-tailed Mann–Whitney test with Prism (GraphPad Software, La Jolla, CA, USA) was also employed for calculating p-value for the distribution of differential expression genes associated to differential accessibility regions. *p*-value and distance (D, maximal vertical distance between the compared distribution) from the comparison of cumulative distribution of RNA levels at 6 hpf were calculated using Kolmogorov-Smirnov Tests by dgof (v 1.4) in R package.

## Data Availability

Raw data are available from corresponding authors upon reasonable request.

## Acknowledgements

We thank all members of the Moreno-Mateos and the Bazzini laboratories for intellectual and technical support. This work was supported by Ramon y Cajal (RyC-2017-23041), PGC2018-097260-B-I00, PID2021-127535NB-I00, MDM-2016-0687 and CEX2020-001088-M grants funded by MICIU/AEI/10.13039/501100011033 by “ERDF A way of making Europe”, “ERDF/EU” and by ESF Investing in your future from Ministerio de Ciencia, Innovación y Universidades and European Union (M.A.M.-M.). M.A.M.-M. was the recipient of the Genome Engineer Innovation 2019 Grant from Synthego. Moreno-Mateos lab has been also co-financed by the Spanish Ministry of Science and Innovation with funds from the European Union NextGenerationEU (PRTR-C17.I1) and the Regional Ministry of University, Research and Innovation of the Autonomous Community of Andalusia within the framework of the Biotechnology Plan applied to Health and grant CNS2022-135564 funded by MICIU/AEI/10.13039/501100011033 and by the European Union NextGenerationEU/PRTR. The CABD is an institution funded by Pablo de Olavide University, Consejo Superior de Investigaciones Científicas (CSIC), and Junta de Andalucía. L.H-H. was a recipient of ayudas para contratos predoctorales para la formación de doctores contemplada en el Subprograma Estatal de Formación del Programa Estatal de Promoción del Talento y su Empleabilidad en I+D+i, en el marco del Plan Estatal de Investigación Científica y Técnica y de Innovación 2017-2020 (Ministerio de Ciencia e Innovación). J.C.C. is supported by Junta de Andalucía Predoctoral Grant (PREDOC_01569) respectively. I.M-S. was a recipient of the Margarita Salas Postdoctoral contract funded by “NextGenerationEU”, Plan de Recuperación, Transformación y Resilencia and Ministerio de Universidades (recualificación del sistema universitario español 2021-2023, Pablo de Olavide University Call). J.M.S-P. was funded by an Emergia grant from Junta de Andalucía (EMC21_00188). This study was supported by the Stowers Institute for Medical Research. A.A.B. was awarded a US National Institutes of Health grant (NIH-R01 GM136849 and NIH R21OD034161). This work was performed as part of thesis research for G.S.P, Graduate School of the Stowers Institute for Medical Research grants. We thank our colleagues Juan Ramon Martínez-Morales, Barbara Pernaute and Thomas Spruce (CABD, Seville, Spain) and Jose Carlos Reyes (CABIMER, Seville, Spain) for the critical reading of the manuscript. We also thanks Rhonda Egidy and Anoja Perera and Ying Zhang and Zhihui (Tim) Wen from Sequencing and Discovery Genomics and Systems Mass Spectrometry facilities at (Kansas, MO, USA), respectively, Danielson Baia Amaral (Stowers Institute) for technical support in performing injections, embryo collection, photography and qPCR for Valine quantification experiments, applying Slam-dunk pipeline and providing suggestions on the manuscript draft and Juan J. Tena (CABD, Seville, Spain) for H3k27ac antibody.

## Author contributions

M.A.M-M. and A.A.B. conceived the project and designed the research. L.H-H. performed all zebrafish experiments with the contribution of I.M-S., A.V-B. and M.A.M-M. J.C-C. carried out medaka experiments. G.dS.P. and J.M.S-P. contributed to the analysis of phospho-proteomic and ATAC-seq analysis. G.dS.P. and L.H-H. performed BCAA quantification experiments. M.A.M-M. and L.H-H. performed data analysis with the help of A.A.B. M.A.M-M. wrote the manuscript with the contribution of L.H-H. and A.A.B. and with the input from the other authors. All authors reviewed and approved the manuscript.

