## Supplementary figures for "CRISPR-RfxCas13d screening uncovers Bckdk as a post-translational regulator of the maternal-to-zygotic transition in teleosts"

##### Supplementary Figure 1. CRISPR-RfxCas13d maternal screening identifies candidates with a role along and after MZT in zebrafish embryos.

**A)** Stacked barplots showing the percentage of wild type (WT), developmentally altered or dead zebrafish embryos at 24 hpf after the depletion of *bckdk*, *cab39l*, *calm1a*, *calm2a*, *mknk1*, *mknk2a* and *ppp4r2a* mRNAs. Embryos were previously divided at 6 hpf according to their developmental phenotype. No: embryos between germ ring and shield stage at 6 hpf. Yes: embryos between 30-50% epiboly at 6 hpf. The results are shown as the averages ± standard error of the mean of each developmental stage from two independent experiments. mRNAs encoding for kinases are labelled in blue and mRNAs encoding for phosphatases are labelled in green. The number of embryos evaluated (n) for each condition is shown.

**B)** Stacked barplots showing the percentage of wild type and developmentally altered embryos from the CRISPR-RfxCas13d screening conditions that did not present more than 35% of embryos with epiboly defects at 6 hpf (Fig. 1C). The results are shown as the averages ± standard error of the mean of each developmental stage from at least two independent experiments. mRNAs encoding for kinases are labelled in blue and mRNAs encoding for phosphatases are labelled in green. The number of embryos evaluated (n) for each condition is shown.

**C)** Representative pictures of embryos injected with RfxCas13d and three gRNAs targeting *mibp* mRNA (gMIBP) compared to uninjected embryos (WT) evaluated at 30 hpf (scale bar, 0.5 mm). Class I: curved tail (mild phenotype). Class II: shorter tail and partial microcephaly (severe phenotype). Class III: notochord malformation and microcephaly (extremely severe phenotype).

**D)** Stacked barplots showing percentage of observed phenotypes at 30 hpf after the injection of RfxCas13d protein (Cas13d) (3 ng/embryo) alone or together with a mix of 3 gRNAs targeting *mibp* mRNA (gMIBP) (1000 pg/embryo). The results are shown as the averages ± standard error of the mean of each developmental stage from three independent experiments. The phenotype selection follows the criteria of Supp Fig 1C. The number of embryos evaluated (n) for each condition is shown.

**E)** RT-qPCR analysis showing levels of *mibp* mRNA at 4 hpf in indicated conditions. Results are shown as the averages ± standard error of the mean from 2 experiments with 2 biological replicates per experiment (n = 10 embryos/biological replicate) for RfxCas13d protein alone (Cas13d) and RfxCas13d plus a mix of 3 gRNAs targeting *mibp* mRNA (gMIBP). *taf15* mRNA was used as normalization control (p *< 0.05, unpaired t-test).

**F)** Intersection visualization analysis showing the number of downregulated genes shared between the mRNA knockdowns of *calm1a*, *calm2a* and *cab39l*.

**Supplementary Figure 2. Transcriptome analysis showed a global downregulation of PZG upon *bckdk* and *mknk2a* mRNA depletion.**

Scatter plots representing the fold change in mRNA levels and *p*-value from a minimum of two biological RNA-seq replicates at 4 hpf of 7 candidates with epiboly defects (positive candidate) in at least 35% of injected embryos and 7 candidates that did not pass this developmental phenotype filter (negative candidate). Pure Zygotic Genes mRNAs defined by Lee *et al.,* 2013[29] are indicated in blue. The mRNA targeted in each condition is highlighted in red.

**Supplementary Figure 3. CRISPR-RfxCas13d maternal screening upon our optimized conditions did not show collateral effect.**

RNA integrity analysis (RIN) from samples used for RNA-Seq in Supp. Fig. 2 analysed by Agilent Bioanalyzer 2100. Electrophoresis gel (A) and RIN associated (B). Developmental phenotype associated to each KD condition is indicated as purple dots (positive candidate) or orange dots (negative candidate). RfxCas13d control condition is indicated as grey dots. One-way ANOVA comparing RIN from all the samples between them is shown in B (ns= non-significant).

**Supplementary Figure 4. The developmental phenotype upon *bckdk* maternal mRNA depletion is recapitulated using independent gRNAs in different teleost models.**

**A)** Schematic representation of endogenous *bckdk* mRNA and individual gRNAs employed (Top). RT-qPCR analysis showing levels of *bckdk* mRNA at 4 hpf in zebrafish embryos co-injected using individual gRNAs targeting *bckdk* mRNA and RfxCas13d protein. Results are shown as the averages ± standard error of the mean from two independent experiments with four biological replicates each (n= 10 embryos/ biological replicate). *taf15* mRNA was used as normalization control (****p< 0.0001, one-way ANOVA) (Bottom).

**B)** Stacked barplots showing the percentage of zebrafish embryos in different developmental stages quantified at 6 hpf using individual gRNAs (1000 pg/embryo) targeting *bckdk* mRNA co-injected with RfxCas13d protein (3 ng/embryo). The phenotype selection criteria were the same as described in Fig. 1C. The results are shown as the averages ± standard error of the mean of each developmental stage from at least two independent experiments. Number of embryos evaluated (n) is shown for each condition.

**C)** Stacked barplots showing developmental phenotypes at 6 hpf upon depletion of *bckdk* mRNA using a gRNA targeting the 3'UTR (gBCKDK-UTR-1). The phenotype selection criteria were the same as described in Fig. 1C. The results are shown as the averages ± standard error of the mean of each developmental stage from two independent experiments. Number of embryos evaluated (n) is shown for each condition.

**D)** Stacked barplots showing developmental phenotypes at 6 hpf in zebrafish embryos injected with the cognate *bckdk* mRNA at 10 or 50 pg per embryo or with a mRNA encoding for GFP at 50 pg per embryo. The phenotype selection criteria were the same as described in Fig. 1C. The results are shown as the averages ± standard error of the mean of each developmental stage from two independent experiments. Number of embryos evaluated (n) is shown for each condition.

**E)** RT-qPCR analysis showing levels of *bckdk* mRNA at 4 hpf in zebrafish embryos in the rescue experiment (Fig. 2A). Results are shown as the averages ± standard error of the mean from one to three independent experiments with at least two biological replicates each (n=10 embryos/ biological replicate). *taf15* mRNA was used as normalization control (p ****< 0.0001, unpaired t-test).

**F)** Western blot showing Bckdk-HA protein expression at 6 hpf in zebrafish embryos injected with 10 or 50 pg/embryo of the cognate *bckdk-HA* mRNA (Top panel). Bottom panel shows Stain-Free signal[104] of the gel as loading control.

**G)** Barplots showing GFP fluorescence signal in zebrafish embryos injected with RfxCas13d alone (Cas13d) or with 2 gRNAs targeting *bckdk* mRNA in the coding sequence (Bckdk KD; gBCKDK-2-3) or one in the 3'UTR (Bckdk KD; gBCKDK-UTR-1) together with 50 pg of *gfp* mRNA. GFP signal is quantified from 3 biological replicates of 5 embryos each (ns= non-significant, unpaired t-test). Representative fluorescence microscopy images used for the quantification are shown.

**H)** Stacked barplots representing the percentage of embryos normally developed (WT) or delayed at 2 or 4 hpf in zebrafish embryos injected with RfxCas13d protein alone (Cas13d) (3 ng/embryo) or with a mix of 3 gRNAs (1000 pg/embryo) targeting *bckdk* mRNA (Bckdk KD). The results are shown as the averages ± standard error of the mean of each developmental stage from two independent experiments. Number of embryos evaluated (n) is shown for each condition.

**I)** Number of total cells or percentage of mitotic cells in zebrafish embryos at 4 hpf injected with RfxCas13d protein alone (Cas13d) or with a mix of 3 gRNAs targeting *bckdk* mRNA (Bckdk KD). Black line represents the median from two independent experiments with at least four embryos each. (ns= non-significant, unpaired t-test).

**J)** Representative immunofluorescence used for quantification in panel **I** showing zebrafish embryos injected with RfxCas13d alone (Cas13d) or with gRNAs targeting *bckdk* mRNA (Bckdk KD), fixed at 4 hpf with PFA 4% and incubated with Phalloidin and DAPI (see Methods for more details). (Scale bar, 100 µm).

**K)** Stacked barplots representing the percentage at 48 hpf of wild type (WT), developmentally altered (DA) or dead medaka embryos injected with RfxCas13d protein (6 ng/embryo) alone or together with a gRNA (gBCKDK-Med) (2000 pg/embryo) targeting *bckdk* mRNA. The results are shown as the averages ± standard error of the mean of each developmental stage from three independent experiments. Number of embryos evaluated (n) is shown for each condition.

**L)** Scatter plots representing the fold change in mRNA level and the associated *p*-value from three biological RNA-seq replicates (n=10 embryos/biological replicate) at 8 hpf in medaka embryos from the comparison between embryos injected only with RfxCas13d protein or together with a gRNA targeting *bckdk* mRNA. *bckdk* mRNA is represented in red. Pure Zygotic Genes mRNAs (PZG) from medaka embryos determined by Li *et al.,* 2020[57] data are depicted in blue.

**Supplementary Figure 5. Transcriptomic and genome landscape analysis in *bckdk* knockdown conditions.**

**A)** Stacked barplots showing developmental phenotypes at 6 hpf in zebrafish embryos injected with RfxCas13d protein (3 ng/embryo) or *RfxCas13d* mRNA (150 pg/embryo) alone or together with different amount of $S_4U$ (25mM, 50mM or 75mM). The phenotype selection criteria were the same as described in Fig. 1C. The results are shown as the averages ± standard error of the mean of each developmental stage from at least two independent experiments. Number of embryos evaluated (n) is shown for each condition.

**B)** Scatter plot representing the fold change in mRNA level from unlabeled reads (SLAM-Seq data) and the associated *p*-value from 2 and 4 biological replicates (n= 25 embryos/biological replicate) at 4 hpf from embryos injected with RfxCas13d protein alone or with a mix of 2 gRNAs targeting *bckdk* mRNA, respectively. *bckdk* mRNA is represented in red. Pure Zygotic Genes mRNAs (PZG) determined by Lee *et al.,* 2013[29] data are depicted in blue.

Scatter plots representing the fold change in mRNA level from labeled reads and the associated *p*-value from three biological RNA-seq replicates of zebrafish embryos at 4 hpf from 2 and 4 biological replicates (n= 25 embryos/biological replicate) from embryos injected with RfxCas13d protein alone or with a mix of 2 gRNAs targeting *bckdk* mRNA, respectively. *bckdk* mRNA is represented in red. Pure Zygotic Genes mRNAs (PZG) determined by Lee *et al.,* 2013[29] data are depicted in blue in panel **C** and an update list of Pure Zygotic Genes mRNAs (PZG) determined by Baia-Amaral et al., 2024[6] data are depicted in blue in panel **D**.

**E)** Stacked barplot showing percentage of up-regulated genes in SLAM-Seq data upon *bckdk* mRNA depletion that belong to different categories. Genes were classified according to Baia-Amaral *et al*., 2024[6]. No MZT: Genes that are not present between 0-7 hpf; Maternal: Genes maternally provided as mRNA but not zygotically transcribed; Maternal-and-Zygotic: Genes maternally provided and zygotically transcribe between 4-7 hpf; or Pure-Zygotic: Genes not maternally provided as mRNA and zygotically transcribed between 4-7 hpf.

**F)** Stacked barplot showing percentage of up-regulated MZT genes (Maternal-and-Zygotic or Pure-Zygotic genes) in SLAM-Seq data upon *bckdk* mRNA depletion that belong to different categories. Genes were classified according to transcript levels in wild type conditions at 4 hpf in labelled data6 .Higher transcribed: Genes with more than 10 CPM (Counts per million) in labelled data at 4 hpf in WT conditions; Earlier transcribed: Genes with less than 10 CPM in labelled data at 4 hpf in WT conditions.

**G)** Gene Ontology enrichment analyses of biological processes for down-(Down) or up-regulated genes (Up) from the comparison of SLAM-Seq data between zebrafish embryos injected with RfxCas13d alone and together with 2 gRNAs targeting *bckdk* mRNA. Terms with a false discovery rate (FDR) lower than 0.05 and with more than 20 genes represented are shown and considered as enriched.

**H)** Differential analyses of chromatin accessibility between zebrafish embryos injected with RfxCas13d protein alone or co-injected with 2 gRNAs targeting *bckdk* mRNA from ATAC-seq data at 4 hpf from 2 biological replicates per condition (n= 80 embryos/biological replicate). The Log2 fold change (FC) and *p*-value associated is represented. Regions with a significant decrease in accessibility are represented as purple dots (log2FC < -1 and *p*-value < 0.05) and regions with a significantly increase in accessibility are represented as orange dots (log2FC > 1 and *p*-value < 0.05). Vertical and horizontal dashed lines indicate 1.5-fold and *p*-value = 0.05, respectively.

**I)** Motif enrichment analyses from the decreased ATAC regions (down peaks) in *bckdk* mRNA knockdown condition. Top-5 motifs are represented with their motif logos, Transcription Factor name, percentage of peaks containing the motif and enrichment *p*-value.

**J)** Motif enrichment analyses from the increased ATAC regions (up peaks) in the same conditions than panel **I**. Top-5 motifs are similarly represented as in panel **I**.

**K)** Stain-Free signal[105] of the gels employed as loading control for Western-Blot in Fig. 3F.

**Supplementary Figure 6. *bckdk* mRNA depletion affects the processing of *miR-430***

**A)** Schematic representation of miR-430 triplet, RT-qPCR primer employed for measuring of primary *miR-430* levels are indicated with black arrows (Adapted from Hadzhiev et al., 2023[94]).

**B)** RT-qPCR analysis showing levels of *mature miR-430* isoforms (*miR430-a*, red; *miR430-b*, orange; and *miR430c*, yellow) at 4.3 hpf (Dome) and 6 hpf (Shield). Results are shown as the averages ± standard error of the mean from 3 experiments with 2 biological replicates per experiment (n= 10 embryos/biological replicate) for RfxCas13d protein alone and RfxCas13d plus 2 gRNAs targeting *bckdk* mRNA. ncRNA *u4atac* was used as normalization control (ns= non-significant, *p<0.05, **p<0.01, unpaired t-test). Same data as represented in Fig. 4C.

**Supplementary Figure 7. Phospho-proteomic analysis upon *bckdk* mRNA depletion identified several potential proteins controlling MZT.**

Schematic representation of the proteins found less **(A)** or more **(B)** phosphorylated upon *bckdk* mRNA knockdown condition. Its known cell localization[106] and the target residues are represented.

**C)** Scatter plot representing the fold change in protein level and the associated *p*-value from 4 biological replicates (n= 100 embryos/biological replicate) at 4 hpf from embryos injected with RfxCas13d protein alone or with a mix of 2 gRNAs targeting *bckdk* mRNA, respectively. Less (Down) or more (Up) abundant proteins are indicated in purple and yellow, respectively. Dashed lines indicated 2-fold in proteins levels and *p*-value= 0.05.

**D)** Stacked barplots showing the percentage of phenotypes observed at 6 hpf from embryos injected with RfxCas13d (3 ng/embryo) alone or together with indicated gRNAs (1000 pg/embryo) targeting *phf10* mRNA. The results are shown as the averages ± standard error of the mean of each developmental stage from at least two independent experiments. The phenotype selection criteria were the same as in Fig. 1C. Number of embryos evaluated (n) is shown for each condition.

**E)** RT-qPCR analysis showing levels of *phf10* mRNA at 2 hpf in zebrafish embryos co-injected using a gRNA (gPHF10-1) targeting *phf10* mRNA and RfxCas13d protein. Results are shown as the averages ± standard error of the mean from two independent experiments with two biological replicates each (n= 10 embryos/biological replicate). *taf15* mRNA was used as normalization control (****p< 0.0001, unpaired t-test).

**F)** Stacked barplots showing the percentage of phenotypes observed at 6 hpf from embryos injected with 50 pg/embryo of *phf10* mRNA WT (Phf10-16T) or modified in the residue phosphorylated by BCKDK. Phf10-16D (Aspartic acid, mimic the phosphorylation state mediated by Bckdk), Phf10-16A (Alanine, mimic a constitutively non-phosphorylated version). The number of embryos evaluated (n) for each condition is shown. The results are shown as the averages ± standard error of the mean of each developmental stage from two independent experiments. The phenotype selection criteria were the same as in Fig. 1C. Number of embryos evaluated (n) is shown for each condition.

**G)** Western blot showing Phf10-HA expression at 6 hpf in zebrafish embryos injected with 50 pg/embryo of the *phf10-HA* mRNA versions used in **H**. Stain-Free signal[105] of the gel as loading control.

Hernández-Huertas *et al.* Supp. Fig. 1

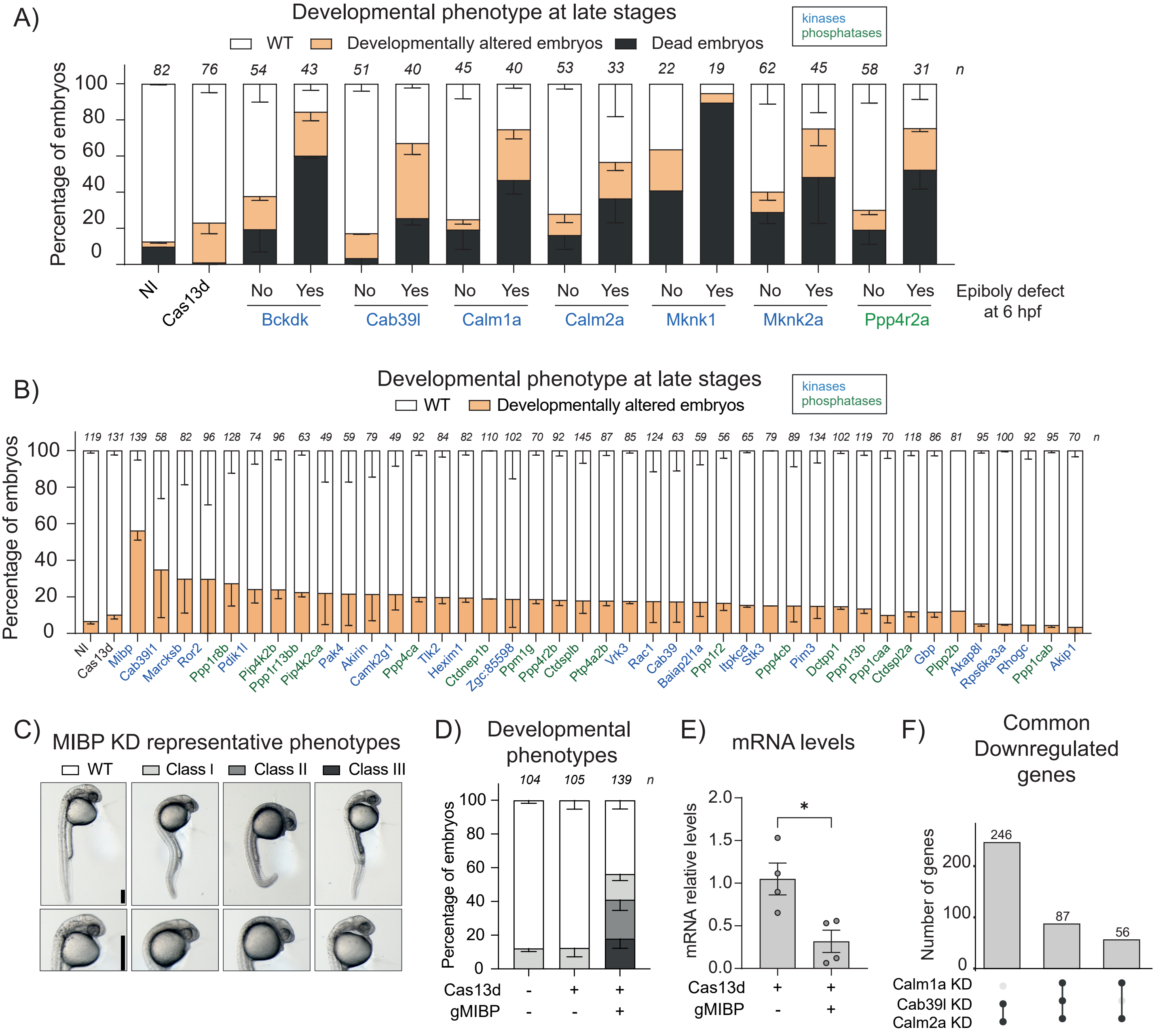

Hernandez-Huertas *et al.*, Supp. Fig. 2

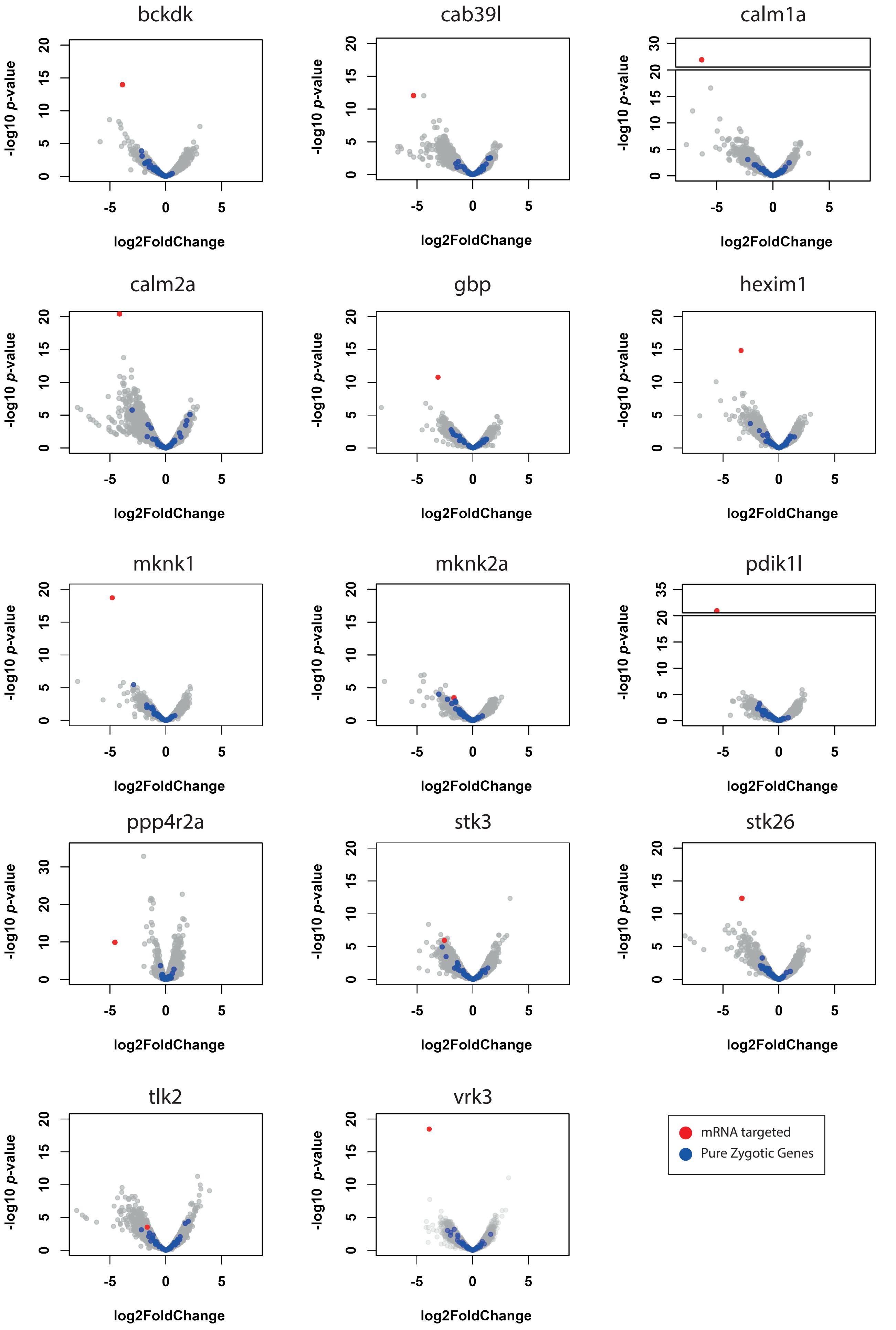

Hernandez-Huertas *et al.*, Supp. Fig. 3

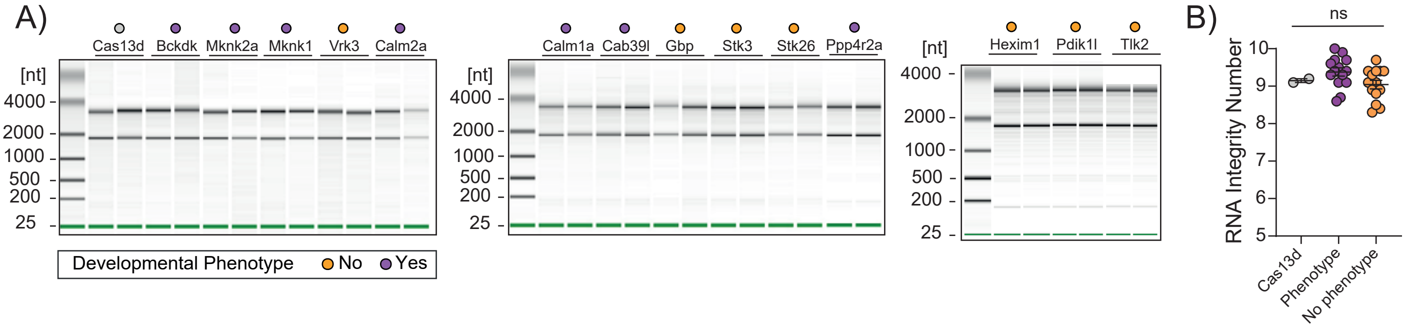

### Hernández-Huertas *et al.* Supp. Fig. 4

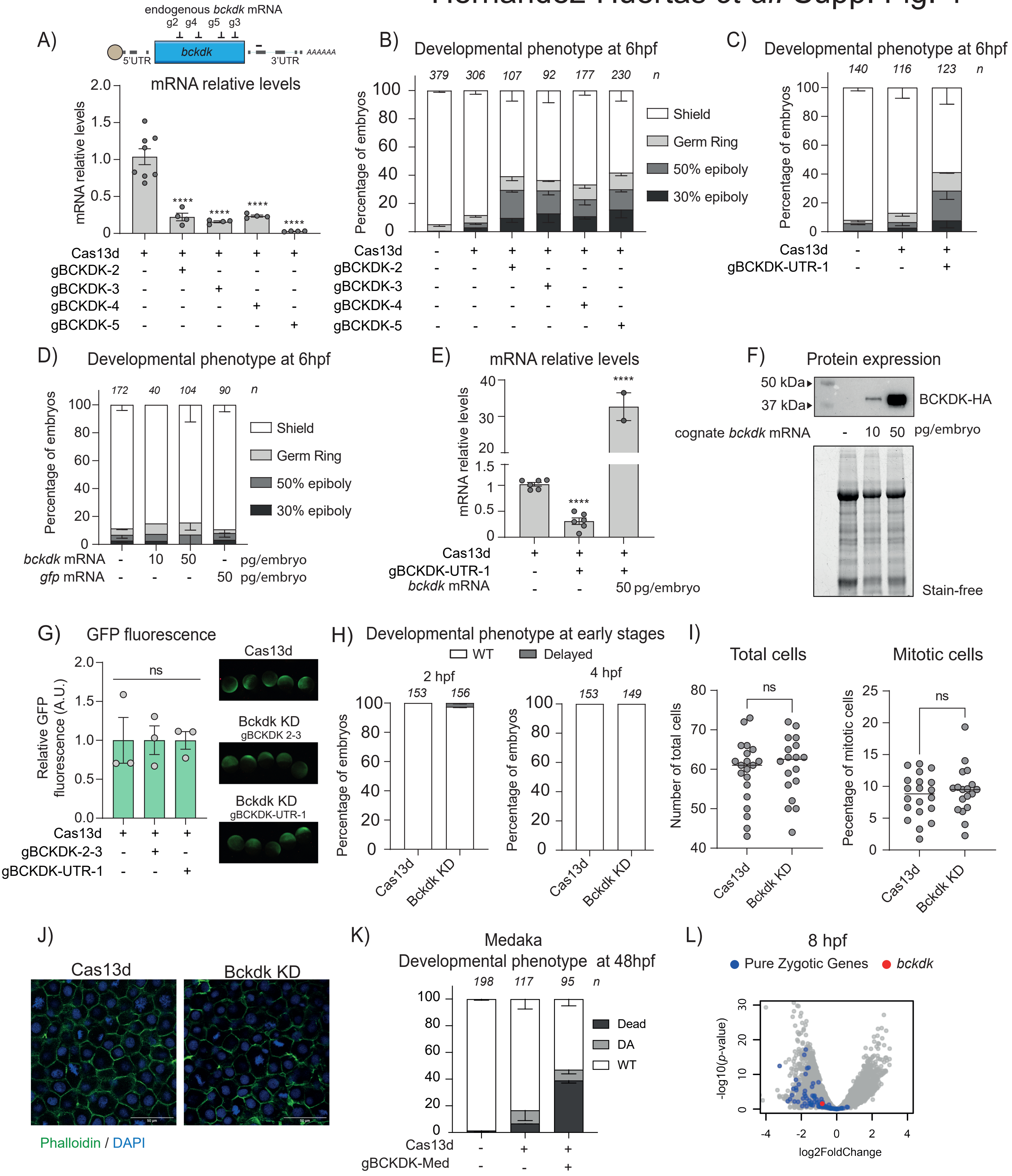

### Hernández-Huertas *et al.* Supp Fig 5

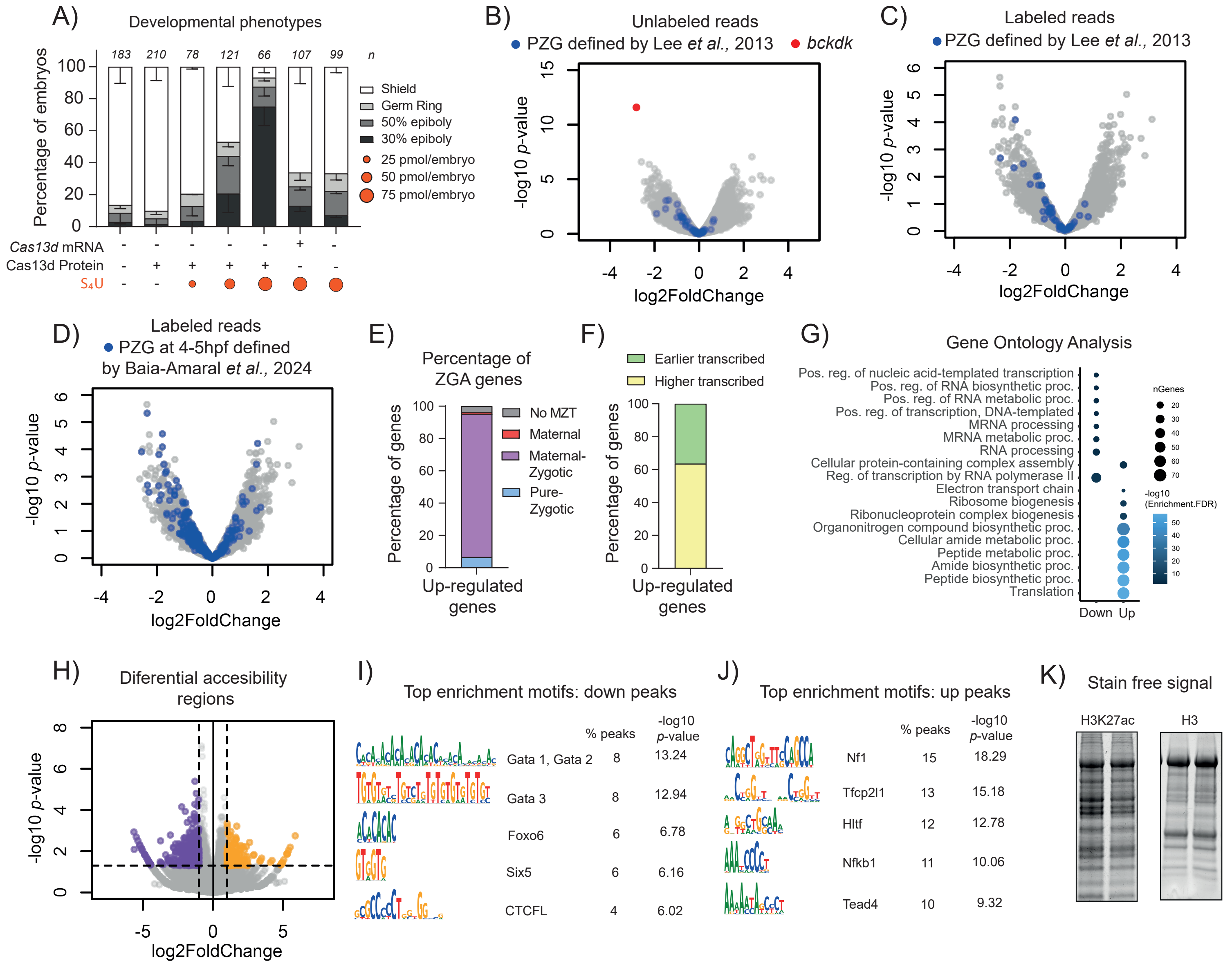

### Hernández-Huertas *et al.* Supp Fig 6

A)

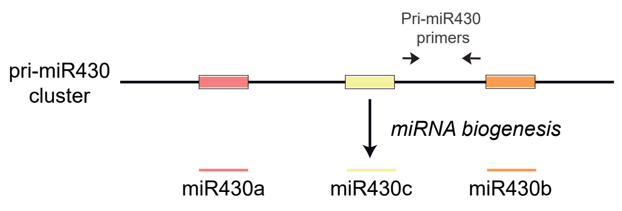


B)

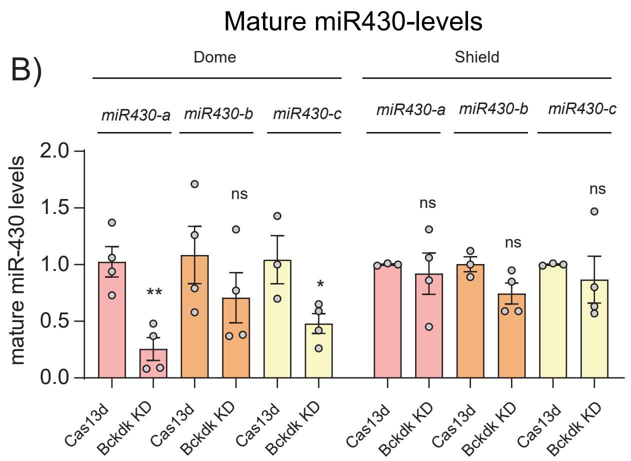

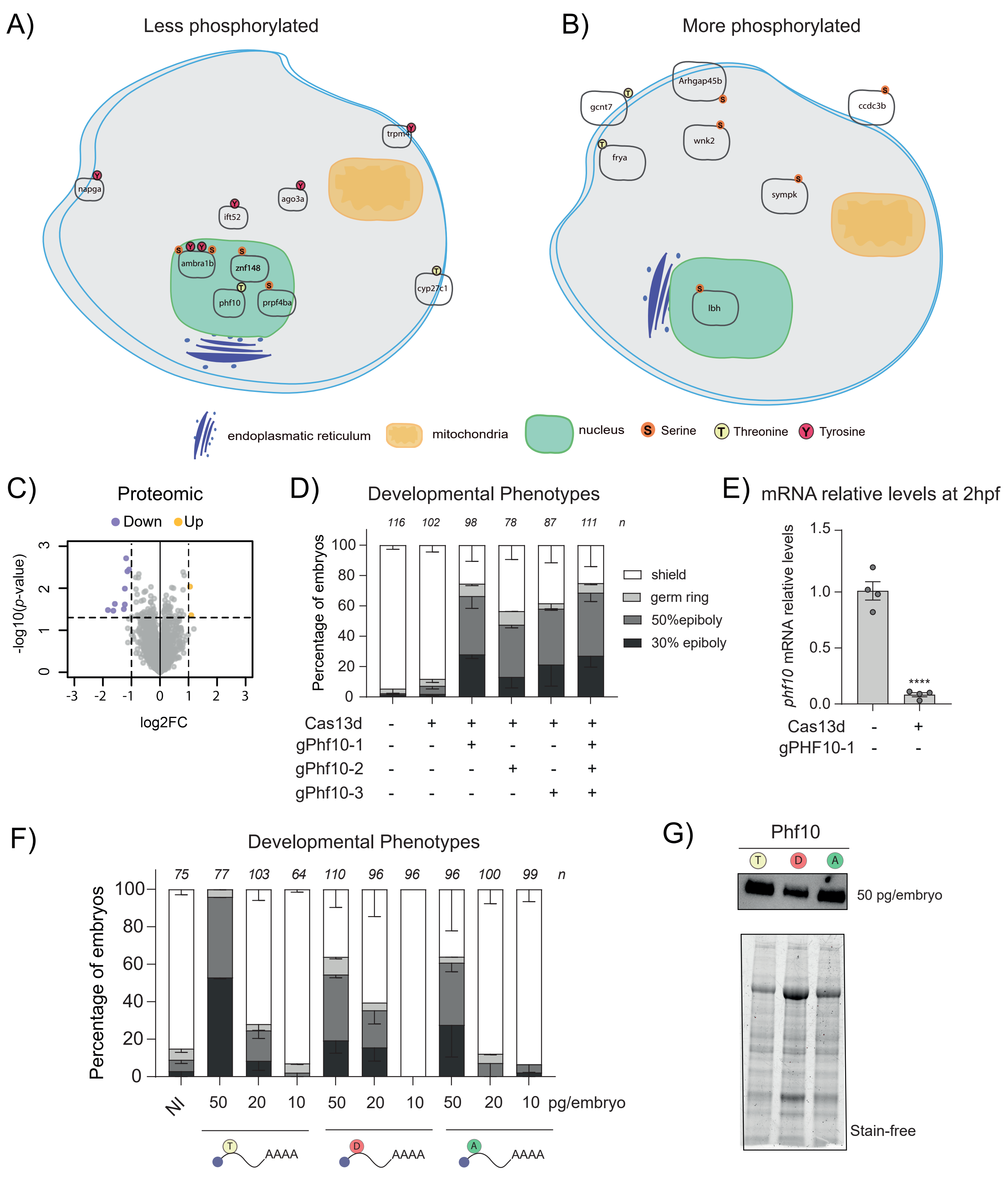

A)
Less phosphorylated
napga
trpm4
ago3a
ift52
ambra1b
znf148
phf10
prpf4ba
cyp27c1
B)
More phosphorylated
gcnt7
Arhgap45b
ccdc3b
wnk2
frya
sympk
lbh
endoplasmatic reticulum
mitochondria
nucleus
Serine
Threonine
Tyrosine
C)
Proteomic
Down
Up
-log10(p-value)
log2FC
D)
Developmental Phenotypes
Percentage of embryos
116 102 98 78 87 111 n
shield
germ ring
50%epiboly
30% epiboly
Cas13d
gPhf10-1
gPhf10-2
gPhf10-3
E)
mRNA relative levels at 2hpf
phf10 mRNA relative levels
****
Cas13d
gPHF10-1
F)
Developmental Phenotypes
Percentage of embryos
75 77 103 64 110 96 96 96 100 99 n
NI 50 20 10 50 20 10 50 20 10 pg/embryo
AAAA
G)
Phf10
50 pg/embryo
Stain-free
